# Defective membrane repair machinery impairs survival of invasive cancer cells

**DOI:** 10.1101/2020.04.20.050211

**Authors:** F. Bouvet, M. Ros, E. Bonedeau, C. Croissant, L. Frelin, F. Saltel, V. Moreau, A. Bouter

## Abstract

Cancer cells are able to reach distant tissues by migration and invasion processes. Enhanced ability to cope with physical stresses leading to cell membrane damages may offer to cancer cells high survival rate during metastasis. Consequently, down-regulation of the membrane repair machinery may be a therapeutic avenue for inhibiting metastasis. We show that migration of MDA-MB-231 cells on collagen I fibrils induces disruptions of plasma membrane and pullout of membrane fragments in the wake of cells. These cells are able to reseal membrane damages thanks to annexins (Anx) that are highly expressed in invasive cancer cells. *In vitro* membrane repair assays reveal that MDA-MB-231 cells respond heterogeneously to membrane injury and some of them possess very efficient repair machinery. Finally, we show that silencing of AnxA5 and AnxA6 leads to major defect of the membrane repair machinery responsible for the death of migrating MDA-MB-231 cells. Inhibition of membrane repair machinery may therefore represent a promising avenue for annihilating cancer metastasis.

**Summary:** Cancer cells are able to reach distant tissues by migration and invasion processes. This study shows that inhibition of the plasma membrane repair machinery may represent a promising avenue for annihilating cancer metastasis.

## Introduction

In cells exposed to mechanical stress, such as skeletal or cardiac muscle cells, epithelial cells and endothelial cells, plasma membrane disruption is a physiological event that occurs frequently (McNeil and Ito, 1989, 1990; McNeil and Khakee, 1992; Clarke et al., 1995). These cells possess a molecular machinery enabling repair of membrane disruptions at the minute scale (McNeil and Steinhardt, 2003; Jimenez and Perez, 2017). The absence of membrane resealing leads to cell death and may contribute to the development of degenerative diseases, such as muscular dystrophies (Griffin et al., 2016; Bansal et al., 2003). Various mechanisms enabling membrane repair have been described such as protein clogging, membrane injury endocytosis or shedding, and lipid patching(Jimenez and Perez, 2017). Influx of Ca^2+^ from extracellular (mM-) to intracellular (µM-range concentration) is the main trigger of membrane repair, which relies mainly on Ca^2+^-sensitive membrane binding proteins such as dysferlin (Glover and Brown, 2007), AHNAK (Huang et al., 2007), calpains (Chrobáková et al., 2004) and Anx (Lennon et al., 2003; Bouter et al., 2011; Roostalu and Strähle, 2012; Boye et al., 2017; Koerdt and Gerke, 2017). MG53 is also a crucial component of the membrane repair machinery, which is activated by the change of oxidation state instead of Ca^2+^ increase due to entry of extracellular milieu (Cai et al., 2009). In the case of mechanical tear, the membrane repair machinery conducts the fusion of intracellular vesicles, which builds a lipid “patch”. This lipid “patch” is recruited to the wounded site via an exocytic-like process for resealing the damaged plasma membrane (Terasaki et al., 1997; McNeil et al., 2000; Carmeille et al., 2016).

Anx appear to be crucial components of the membrane repair machinery in a wide range of cell types. The Anx family consists of twelve soluble proteins in mammals, named AnxA1 to A13 (the number 12 is non-assigned) (Moss and Morgan, 2004). In a Ca^2+^-dependent manner, Anx share the property of binding to membranes exposing negatively charged phospholipids (Moss and Morgan, 2004; Gerke et al., 2005). They exhibit little specificity for anionic lipid head-groups and variable threshold-value of Ca^2+^ concentration for membrane binding (Andree et al., 1990; Blackwood and Ernst, 1990; Monastyrskaya et al., 2007). In mouse primary muscle cells, AnxA1 and AnxA2 may be responsible for fusion of intracellular vesicles and recruitment of lipid “patch” to the membrane disruption site by interacting with dysferlin (Lennon et al., 2003). In skeletal muscle of zebrafish, it has been proposed that membrane repair is based on the establishment of a highly ordered scaffold involving dysferlin, AnxA6, AnxA2 and AnxA1 (Roostalu and Strähle, 2012). In the presence of Ca^2+^, AnxA5 presents the property of rapidly assembling into trimers and self-organizing into 2D ordered arrays, when it binds to biological membranes (Kaetzel et al., 2001; Lambert et al., 1997; Oling et al., 2001; Voges et al., 1994). We have shown that AnxA5 promotes membrane repair by forming 2D arrays at the torn membrane edges, thus strengthening the membrane, preventing the expansion of wound and facilitating the final steps of membrane resealing (Bouter et al., 2011; Carmeille et al., 2015, 2016).

As a defect in membrane repair is responsible for the development of muscular dystrophies (Liu et al., 1998; Bansal et al., 2003; Minetti et al., 1998), major attention has focused on membrane repair of skeletal muscle cells, leaving membrane repair in other tissues and in other pathologies almost not characterized. Cancer cell metastasis is a cascade of events that allows the dissemination of the disease from the site of origin into peripheral vessels, and, subsequently, other organs. One of the hallmarks of metastatic cells is their ability to invade the surrounding tissues. Tumor cell migration and invasion require actin cytoskeleton remodeling, which increases membrane dynamics and reduces stiffness of the plasma membrane (Swaminathan et al., 2011). In addition, invasive cancer cells are submitted to tremendous shearing forces, due to their migration through the extracellular milieu and their invasion in distant tissue, which may lead to numerous plasma membrane damages (see for review (Wirtz et al., 2011)).

Even though few studies are linked membrane repair and cancer, numerous works have highlighted incidentally a positive correlation between major actors of the membrane repair machinery and tumor invasion. For instance, it has been observed that the overexpression of many Anx enhances tumor aggressiveness and is a factor of poor prognosis in various cancers such as breast cancer (Chuthapisith et al., 2009; MA et al., 2014; Yang et al., 2013; Yao et al., 2009; Zhang et al., 2012; Wei et al., 2015; Deng et al., 2013; Duncan et al., 2008). In addition, Anx inhibition significantly reduces proliferation, migration and invasion abilities of tumor cells, without knowing accurate mechanism (Wehder et al., 2009; Peng et al., 2016; Ding et al., 2017; Wu et al., 2012; Koumangoye et al., 2013). Credit has to be given to J. Jaiswal and J. Nylandsted for the first investigations describing membrane repair machinery in cancer cells. They show that the AnxA2/S100A11 complex promotes membrane repair by remodeling actin cytoskeleton and facilitating excision of the damaged membrane area in MCF-7 cells (Jaiswal et al., 2014). Since, AnxA4 and AnxA6 have also been shown to participate in membrane repair of MCF-7 cells by inducing respectively membrane curvature from the edges of the torn membrane and constriction force to pull the wound edges together (Boye et al., 2017). Finally, the excision of damaged membrane by ESCRT-mediated shedding requires AnxA7 in MCF-7 cells (Sønder et al., 2019). Altogether, these findings suggest that an efficient membrane repair machinery composed by Anx exists in cancer cells and may facilitate tumor invasion processes. The aim of the present study was to investigate whether silencing of proteins from the membrane repair machinery may impair survival of invasive cancer cells. To address this question, we have analyzed the behavior of MDA-MB-231 cells seeded on glass coverslip coated with type I collagen (collagen I) fibrils. MDA-MB-231 breast tumor cell line exhibits strong capacity of invasiveness *in vitro* and *in vivo* as well (Thompson et al., 1992). These cells have constituted therefore a suitable model for analyzing the effect of invasiveness on cell membrane disruption and repair. Collagen I is the most abundant extracellular matrix protein in vertebrates and is overexpressed in a large number of cancers (Ramaswamy et al., 2003; Gilkes et al., 2013). The effect of collagen I on tumor cells is complex and paradoxical. Initially expected to form a physical barrier for limiting the progression of the tumor, collagen I has proved to be a powerful inducer of linear invadosomes, which in turn activate cell invasion by promoting matrix metalloproteases activity that degrades extracellular matrix (Juin et al., 2012). After having characterized that AnxA5 and AnxA6 belong to the membrane repair machinery of MDA-MB-231 cells, we compared the responses to collagen I-mediated membrane injury of control cells to cells rendered deficient for AnxA5 and/or AnxA6. Our results lead us to propose that membrane repair machinery may be considered as a promising target for annihilating cancer metastasis.

## Results

### Collagen I fibrils favor migration of MDA-MB-231 cells and formation of migrasome

First, the migration of MDA-MB-231 cells plated on glass coverslip coated or not with collagen I was compared through analysis by phase-contrast video-microscopy (**Fig. 1**). Tracking of cells was performed in both conditions by means of the MtrackJ plugging from the ImageJ software (Meijering et al., 2012) and start-end distances were measured. Start-end distance is the Euclidian distance between positions of cell at t0 and tend. It therefore positively correlates with the motility of the cell. In the presence of collagen I cells exhibited elongated shape and migrate lengthwise (**Fig. 1 and video S1**), whereas in the absence they presented mostly rounded shape and spinning movements (**Fig. 1 and video S2**). Mean start-end distance of cells seeded on collagen I was significantly (P-value = 5.5E-11 for student test) higher (103 ± 33 µm) than in the absence of collagen I (13 ± 12 µm) after 2 h of migration. We have therefore concluded that collagen I fibrils favor the migration of MDA-MB-231 cells.

**Figure 1.**
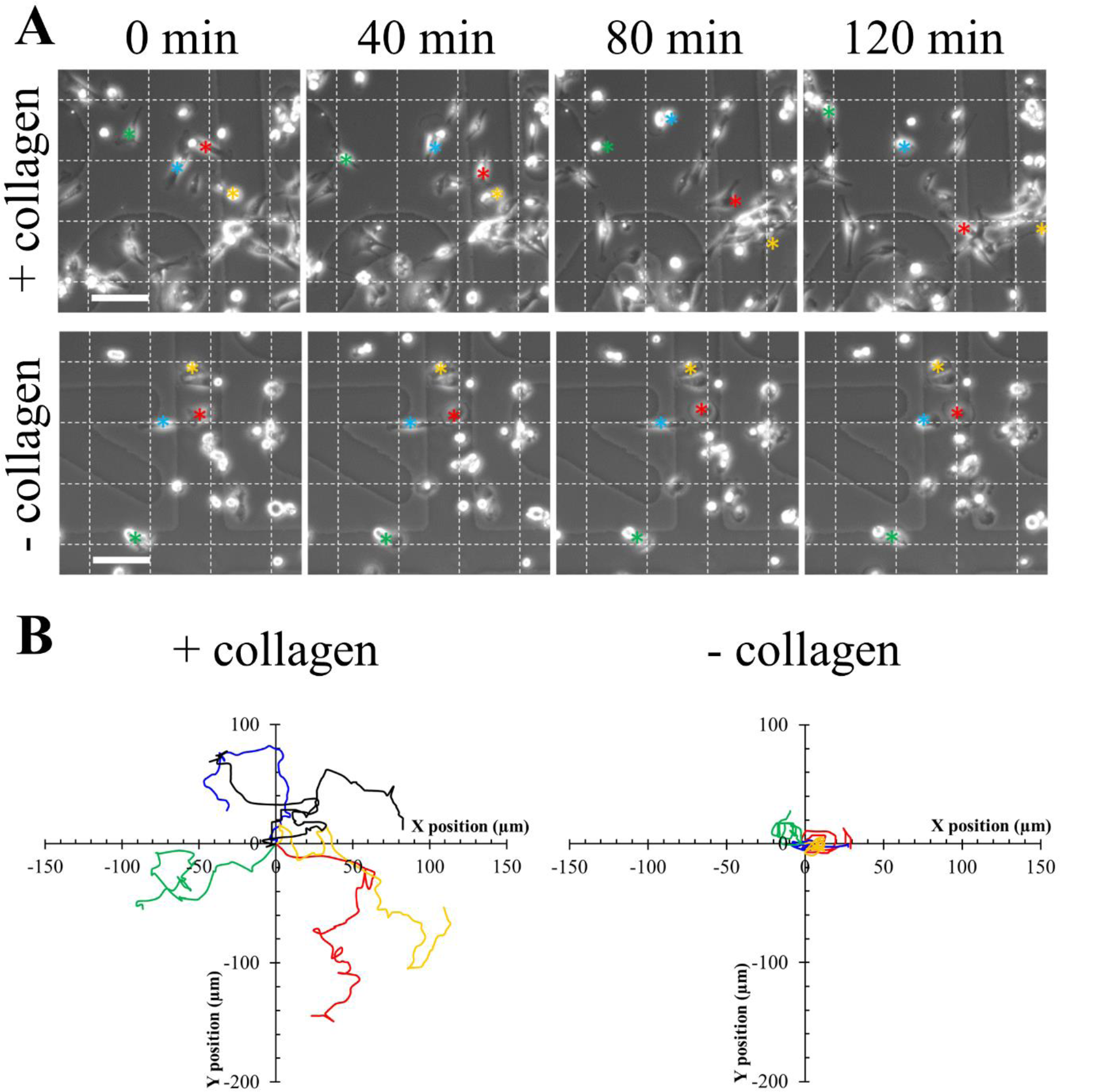
Effect of collagen I fibrils on the migration of MDA-MB-231 cells. **(A)** Kinetics studies by phase-contrast video-microscopy of MDA-MB-231 cells seeded on a glass coverslip coated (top panel) or not (bottom panel) with collagen I. Asterisks (red, blue, green, yellow) correspond to the position of cells of interest in the course of time. Scale bar: 100 µm. **(B)** Tracking analysis of cells presented in (A) was performed using MTrackJ plugging from ImageJ software. Relative position of cells migrating during two hours on glass coverslip coated (left panel) or not (right panel) with collagen I was plotted. For sake of clarity, only four and six tracks were presented in the absence or presence of collagen, respectively. Colors tracks correspond to cells marked with asterisk in (A).

In order to characterize the migrasome (Ma et al., 2015; Wu et al., 2017) of MDA-MB-231 cells moving on collagen I fibrils, cells were cultured on glass coverslip coated with collagen I for 24 h, their migration was analyzed for 2 h by phase-contrast video-microscopy and at the end of the kinetics study cells were fixed and incubated with CellMask™Orange. CellMask™ stains are lipophilic dyes that become fluorescent upon inserting into plasma membrane. The use of glass bottom dishes equipped with a square-patterned coverslip displaying an alphanumerical code in each square enabled cell tracking during different stages of the experiment and correlation of cell migration observed by phase-contrast video-microscopy and CellMask™Orange staining analyzed by fluorescence microscopy. By means of fluorescence microscopy, we observed systematically (over 7 independent experiments) the presence of cell membrane fragments in the near periphery of approximately 70% of cells (**Fig. 2 A**). At higher magnification, cell membrane fragments appeared as membrane-bound vesicular structures (MbVS, **Fig. 2 B**), as previously described by Yu and collaborators (Ma et al., 2015). By analyzing the immediate surroundings of a migrating cell followed by video-microscopy (**Fig. 2 C and video S3**), we observed the presence of MbVS in the wake of the cell by fluorescence microscopy (**Fig. 2 D**). We have concluded that MDA-MB-231 cells migrating on collagen I fibrils release MbVS in their wake. Formation and release of these cell structures may be induced by shearing forces existing between the extracellular collagen I fibrils and cell membrane.

**Figure 2.**
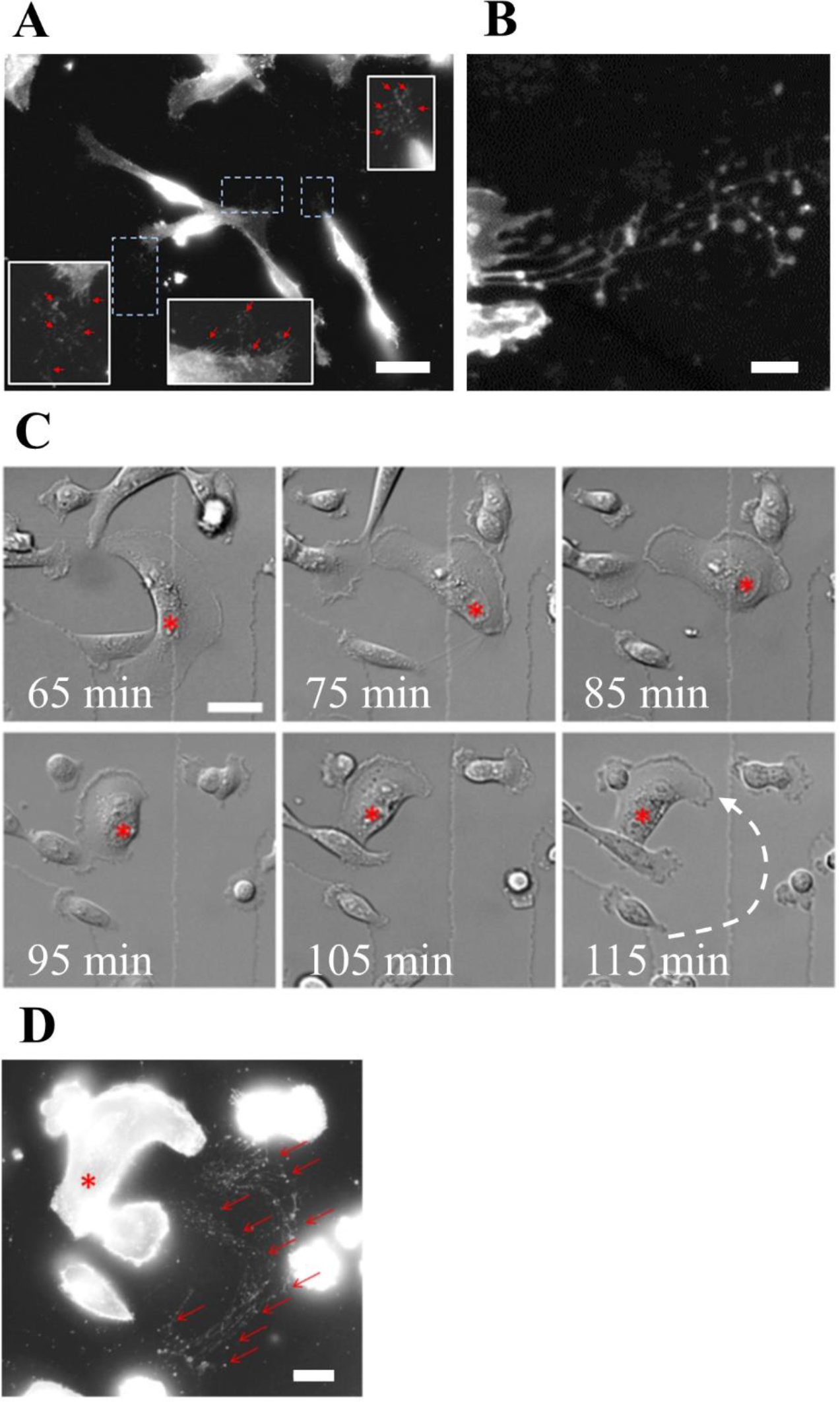
Presence of membrane fragments during MDA-MB-231 cell migration on collagen I fibrils. **(A)** MDA-MB-231 cells were seeded on a glass coverslip coated with collagen I. 24h after seeding, kinetics study of cell migration was performed during 2h by phase-contrast video-microscopy. Cells were then incubated with CellMask™Orange (white). Red arrows indicate membrane material at the periphery of cells. Scale bar: 60 µm. **(B)** Observation at higher-magnification by fluorescence microscopy revealed the presence of membrane material stained by CellMask™Orange (white), which appears as membrane-bound vesicular structures. Scale bar: 5 µm. **(C)** Kinetics study enabled to identify a migrating cell, for which the nucleus has been marked by a red asterisk. The dashed white arrow indicates the path of the cell during the migration. Scale bars: 40 µm. **(D)** After migration, cells were immediately incubated with CellMask™Orange (white) for 5 min and fixed with 4% paraformaldehyde. Red asterisk marks the nucleus of the cell of interest presented in **c** and red arrows indicate membrane fragments present in the wake of the cell. Scale bar: 20 µm.

### Cell membrane disruption and repair in MDA-MB-231 migrating on collagen I

In order to assess if release of MbVS was accompanied by cell membrane disruptions, MDA-MB-231 cells were loaded with Fluo-4-AM and kinetics study of cell migration on collagen I was performed by fluorescence microscopy. Fluo-4-AM is a dye with a fluorescence intensity that considerably varies as a function of intracellular calcium concentration. Weakly fluorescent in intact cells, where the cytoplasmic concentration of calcium is in the µM range, Fluo-4-AM exhibits a strong fluorescence increase when the integrity of plasma membrane is compromised since the calcium concentration increases strongly (mM). Images acquired in phase-contrast microscopy just before and after kinetics study ensured that cells have migrated on collagen I fibrils (**Fig. 3 A**). We observed important variation of Fluo-4-AM fluorescence in several cells during migration (**Fig. 3 B and video S4**). The strong increase of fluorescence intensity revealed the presence of rupture(s) in cell membrane (**Fig. 3, B and C, t=32min or t=85min**). The recovery to a basal fluorescence intensity after about 1-2 min indicated that cell membrane resealed and calcium homeostasis was restored at the minute range (**Fig 3, B and C, t=34min or t=86min**), in line with the timescale required for membrane repair in cells damaged by laser ablation (Bouter et al., 2011; Carmeille et al., 2015, 2016; Koerdt and Gerke, 2017; McNeil and Kirchhausen, 2005). We have concluded that shearing forces between collagen I fibrils and cell membrane of migrating MDA-MB-231 induce membrane disruptions that reseal at the minute range.

**Figure 3.**
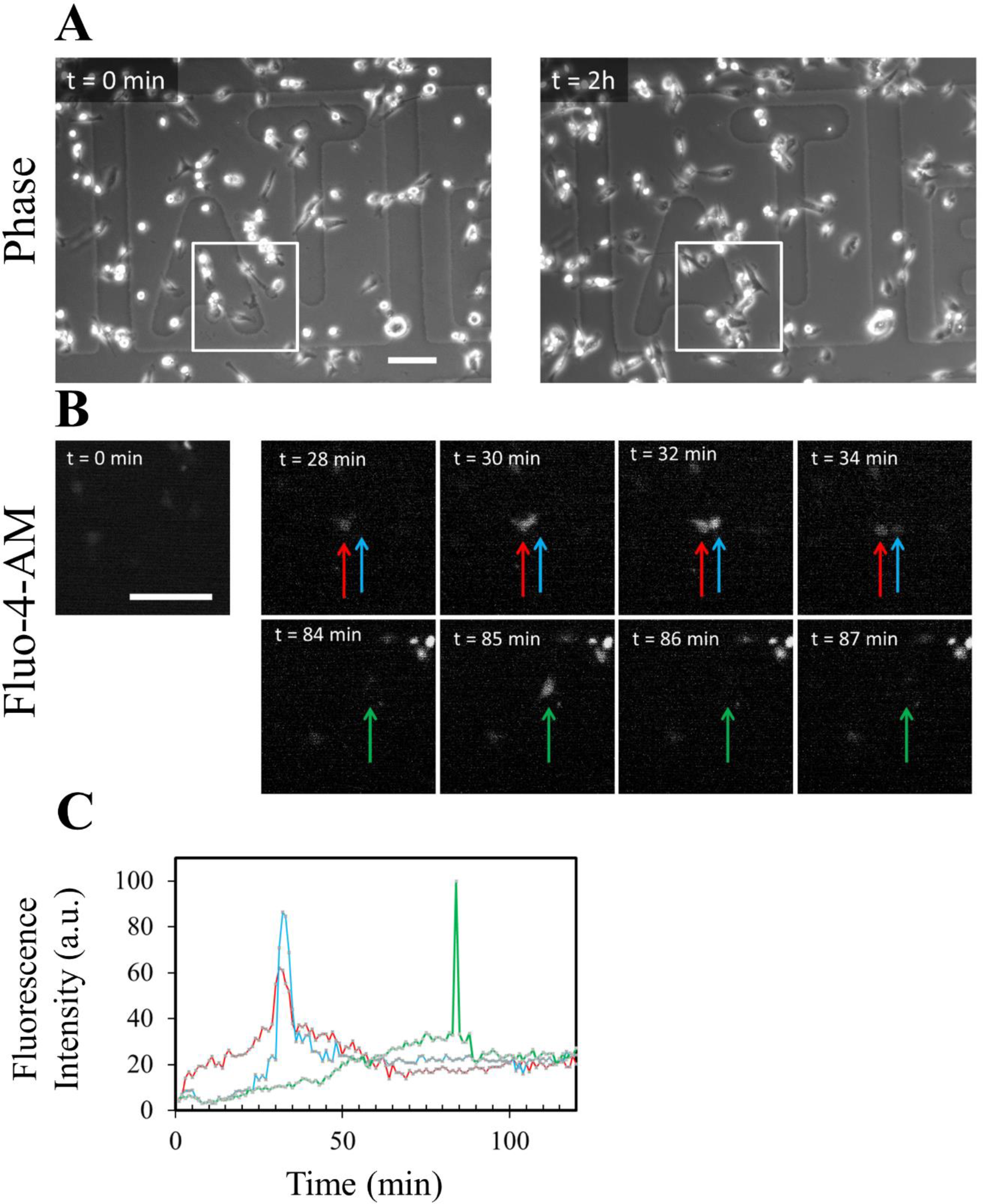
Highlighting membrane ruptures in migrating MDA-MB-231 cells. MDA-MB-231 cells were seeded on a glass coverslip coated with collagen I for 24h, then incubated for 1h at 37°C with 2.5µM Fluo-4-AM (white) and finally kinetics study using fluorescence microscopy was performed for 2h. **(A)** Images in phase-contrast microscopy were acquired before and after the kinetics study. **(B)** Fluorescence intensity variations of intracellular Fluo-4-AM studied in the course of time. Each colored arrow points out a cell of interest exhibiting a strong variation of the fluorescence intensity. A strong increase of fluorescence intensity is particularly observed at t=32min (blue and red arrows) and t=85min (green arrow). A recovery of fluorescence intensity to a basal level is observed at t=34min (blue and red arrows) and t=86min (green arrow). Scale bar: 100 µm. **(C)** Quantification of intracellular fluorescence intensity in arbitrary unit (a.u.) measured in cells presented in (B) during the two hours of migration. The color of each line corresponds to the color of the arrow in (B).

### MDA-MB-231 cells exhibit a highly efficient membrane repair machinery

In order to investigate the membrane repair ability of MDA-MB-231 cells, plasma membrane was damaged by laser ablation and membrane repair was assayed by monitoring the kinetics of cell entry of membrane-impermeable FM1-43 dye, using a standard protocol (Carmeille et al., 2017; Bouter et al., 2011). Membrane resealing leads to a stop in the entry of FM1–43 molecules into the cytosol and therefore to a stop in the increase of the intracellular fluorescence intensity, whereas the absence of membrane resealing leads to a continuous entry of FM1-43 into the cytosol and a continuous and strong increase in the fluorescence intensity. Irradiation conditions were adjusted (110 mW) to cause mild membrane injury to cells, characterized by the entry of FM1-43 into the cytosol as early as a few seconds after laser ablation without the presence of a macroscopic tear at the site of membrane irradiation (**Fig. 4, A and B, frames 2, arrow**). Two typical responses were observed within the 120 s following the membrane damage. 50% of cells exhibited an increase of the fluorescence intensity limited to an area closed to the site of membrane injury (**Fig 4 A and video S5 A**). For these cells, we observed that the fluorescence intensity increased for about 80 s before reaching a plateau (**Fig 4 C, black filled circles**). The presence of the plateau indicated that the entry of FM1-43 was stopped and therefore that cell membrane resealed. The second half of cells presented a massive entry of FM1-43 (**Fig. 4 B and video S5 B**) leading to a continuous and huge increase of the fluorescence intensity (**Fig. 4 C, empty circles**), which indicated the absence of membrane resealing. This significant heterogeneity in the response of MDA-MB-231 cells to laser ablation has never been observed in our previous studies performed with perivascular, placental and muscle cells (Bouter et al., 2011; Carmeille et al., 2015, 2016). It suggests a phenotypic variability within the cell population. For the “repairing” MDA-MB-231 subpopulation, kinetics of FM1-43 entry into damaged cells and variations of the intracellular fluorescence intensity within the 120 s following membrane damage are in line with those observed in human myotubes (Carmeille et al., 2016) and syncytiotrophoblasts (Carmeille et al., 2015). We have therefore concluded that MDA-MB-231 cells possess a cell machinery ensuring efficient repair of membrane damage, while this machinery is not systematically operational.

**Figure 4.**
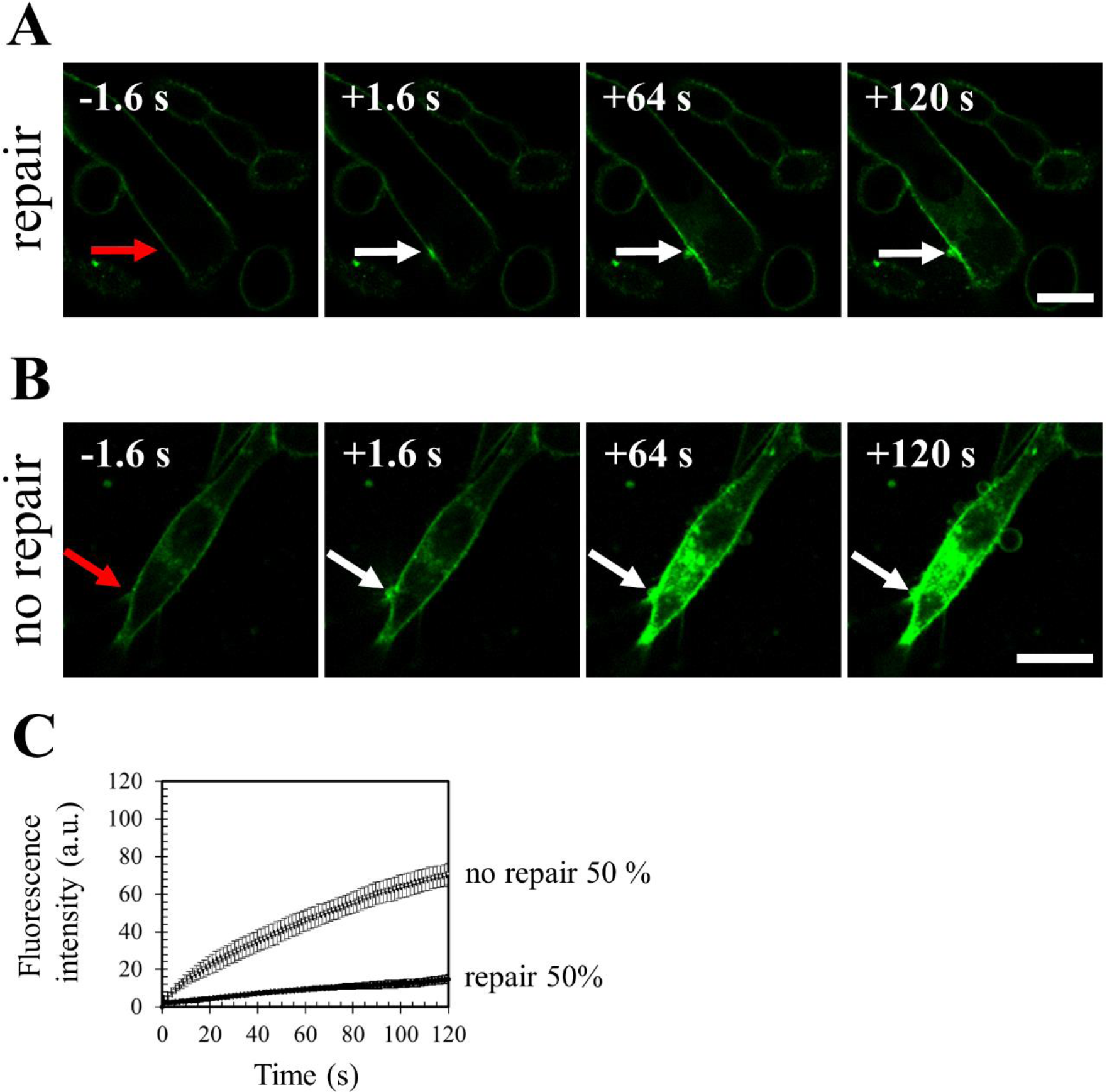
Responses of MDA-MB-231 cells to a membrane damage by laser ablation. Sequence of representative images showing the response of a MDA-MB-231 cell resealing **(A)** or not **(B)** a membrane damage performed by 110-mW infrared laser irradiation, in the presence of FM1-43 (green). Before irradiation, cells were washed 3 times in cold D-PBS containing 2 mM Ca2+ and then maintained in this medium. In all figures, the area of membrane irradiation is marked with a red arrow before irradiation and a white arrow after irradiation. Image frames 1 and 2 were recorded 1.6 s before and 1.6 s after irradiation, respectively; Image frames 3–4 were recorded 64 s and 120 s after irradiation, respectively, as indicated. Scale bar: 20 μm. Representative images of about 100 observed cells. **(C)** Kinetic data represent the FM1−43 fluorescence intensity integrated over whole cell sections, averaged for about 60 cells (+/−SD). For 50% of MDA-MB-231 cells, the fluorescence intensity reached a plateau after about 80 s (black filled circles). For the other half, a continuous and large increase of the fluorescence intensity was observed (empty circles), indicating the absence of membrane resealing.

### Anx are significantly expressed in MDA-MB-231 cells

Several members of the Anx family are instrumental for membrane repair in many cell types, namely AnxA1 (Lennon et al., 2003; Roostalu and Strähle, 2012; Demonbreun et al., 2016), AnxA2 (Jaiswal et al., 2014; Demonbreun et al., 2016; Koerdt and Gerke, 2017), AnxA4 (Boye et al., 2017), AnxA5 (Bouter et al., 2011; Carmeille et al., 2015, 2016) and AnxA6 (Demonbreun et al., 2016; Boye et al., 2017).

In order to assess the relative expression of Anx in MDA-MB-231 cells, we prepared protein extracts from MDA-MB-231 and BeWo cells and analyzed the amount of Anx by western-blot (**Fig. 5, A and B**). The choriocarcinoma BeWo cell line is placental in origin and the placenta is expected to be the richest organ in Anx in Humans (www.proteinatlas.org). BeWo cells served therefore as a positive control of the expression of Anx. In order to standardize measurements, MDA-MB-231 and BeWo protein extracts were analyzed on the same PVDF membrane and the band intensity relative to the Anx was reported to GAPDH, used as loading control. As an example, a typical result obtained for the detection of AnxA5 is presented in **Fig. 5 B**. We observed that AnxA4 is hardly, if at all, expressed in both cell lines. A significant expression of AnxA1, A2, A5, and A6 is observed. For these Anx, the expression level is systematically as high for MDA-MB-231 as for BeWo cells. We even observed that the expression of AnxA2 is stronger in MDA-MB-231 compared to BeWo cells. We can therefore conclude that four Anx (A1, A2, A5 and A6), which are known to participate in membrane repair, are highly expressed in MDA-MB-231 cells.

**Figure 5.**
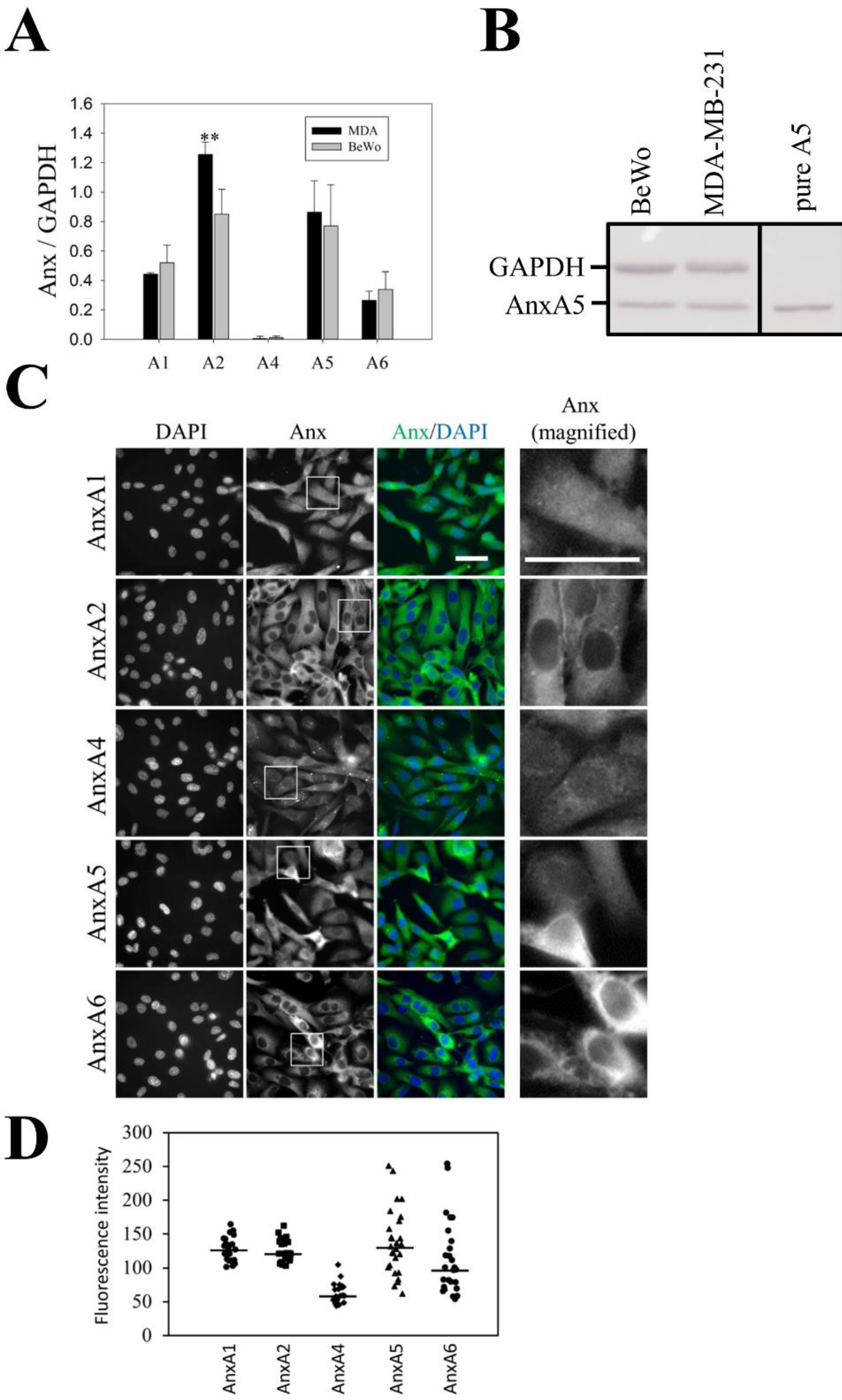
Expression and subcellular localization of endogenous Anx in MDA-MB-231. **(A)** 10 µg of protein extracts from MDA-MB-231 and BeWo cells were analyzed by western-blot. The gel analysis plugging of ImageJ was used to quantify the density of bands. The histogram presents mean values (± SD) of the ratio Anx/GAPDH from 10 experiments. Student t Test for independent samples. **: p < 0.01. **(B)** Revelation of AnxA5 in MDA-MB-231 and BeWo cells compared to 50 ng purified recombinant AnxA5 (Pure A5), presented as an example. **(C)** MDA-MB-231 cells were immunostained for AnxA1, AnxA2, AnxA4, AnxA5 and AnxA6 (green), as indicated, and counterstained with DAPI (blue). The right-hand column presents magnified images extracted from the “Anx” column (indicated by inserts). Scale bar: 40 µm. **(D)** Quantification of fluorescence intensities after immunostaining of endogenous Anx in MDA-MB-231 cells. Fluorescence intensities were measured from 8-bit images by drawing a circular ROI inside the cytoplasm and measuring the mean pixel value. Mean pixel values from at least 20 cells are presented. Horizontal bars represent median values. The variance value is 323, 268, 224, 2372 and 2909 for AnxA1, A2, A4, A5 and A6, respectively. The coefficient of variation is 14%, 13%, 24%, 35% and 47% for AnxA1, A2, A4, A5 and A6, respectively. The expression level of AnxA5 or AnxA6 is heterogenous from one MDA-MB-231 cell to another. In some cells, AnxA5 and AnxA6 are expressed at a very low level.

We then investigated the subcellular distribution of endogenous Anx in MDA-MB-231 cells. Immunocytofluorescence experiments confirmed that AnxA1, AxnA2, AnxA5 and AnxA6 are significantly expressed in MDA-MB-231 cells whereas AnxA4 is weakly expressed (**Fig. 5 C**). As observed in human myoblasts (our personal data) and murine neuroblasts (Skrahina et al., 2008), AnxA1 localizes in the nucleus and in the cytoplasm of MDA-MB-231 cells. Instead AnxA2, AnxA5 and AnxA6 are exclusively cytoplasmic. These results are consistent with previous studies for AnxA2 (Skrahina et al., 2008) and AnxA6 (Vilá de Muga et al., 2009) but different for AnxA5 (Skrahina et al., 2008; Carmeille et al., 2015), which has been reported to be present in the nucleus and the cytoplasm. The homogenous distribution of Anx within the cytoplasm supposes that it localizes in the cytosol. Strikingly, we observed that the expression level of AnxA5 or AnxA6 varies considerably from one MDA-MB-231 cell to another (**Lower panels of Fig. 5, C and D**), whereas that of AnxA1 or AnxA2 is relatively constant (**Higher panels of Fig. 5, C and D**). We observed that a cell expressing AnxA5 at a high level expresses also strongly AnxA6 (**Fig. S1**). In addition, a low level of AnxA5 is systematically accompanied by a weak expression of AnxA6. This strong heterogeneity in the expression of AnxA5 and AnxA6 in MDA-MB-231 cells and the fact that some of them express at a low level both Anx is in line with the absence of membrane repair observed in a part of these cells damaged by laser ablation. This result suggests that AnxA5 and/or AnxA6 may participate in the membrane repair machinery of the MDA-MB-231 cells. Hereafter we have therefore focused our attention on AnxA5 and AnxA6.

### AnxA5 and AnxA6 belong to the membrane repair machinery of MDA-MB-231 cells

Most, if not all, proteins that are involved in the repair machinery, i.e. dysferlin (Bansal et al., 2003), AnxA1, AnxA2, AnxA6 (Lennon et al., 2003; McNeil et al., 2006; Roostalu and Strähle, 2012), AnxA5 (Bouter et al., 2011; Carmeille et al., 2015) and MG-53 (Cai et al., 2009) are recruited to the membrane disruption site immediately after membrane injury. In order to assess whether AnxA5 and AnxA6 may be involved in the membrane repair machinery of MDA-MB-231 cells, we investigated the subcellular localization of both Anx after laser ablation. For this purpose, MDA-MB-231 cells were cultured on gridded coverslips, thus enabling accurate tracking of irradiated cells. After laser irradiation, cells were fixed, permeabilized and immunostained for AnxA5 **(Fig. 6 A)** or AnxA6 **(Fig. 6 B)**. After laser injury, endogenous AnxA5 and AnxA6 were found specifically accumulated at the disruption site of MDA-MB-231 cells. We have previously shown that AnxA5 promotes membrane repair mainly by interacting with the edges of the torn membrane, where it forms bi-dimensional arrays that strengthen the membrane and prevent the expansion of the tear (Bouter et al., 2011; Carmeille et al., 2016). In addition, it has been proposed that AnxA6 is recruited to the wound edges, where it induces a constriction force enabling wound closure (Boye et al., 2017). Interaction between Anx and damaged plasma membrane requires only the presence of phosphatidylserine in the membrane and calcium at mM concentration. Therefore, it is likely that the recruitment of AnxA5 and AnxA6 at the disruption site is responsible for membrane resealing in MDA-MB-231 cells.

**Figure 6.**
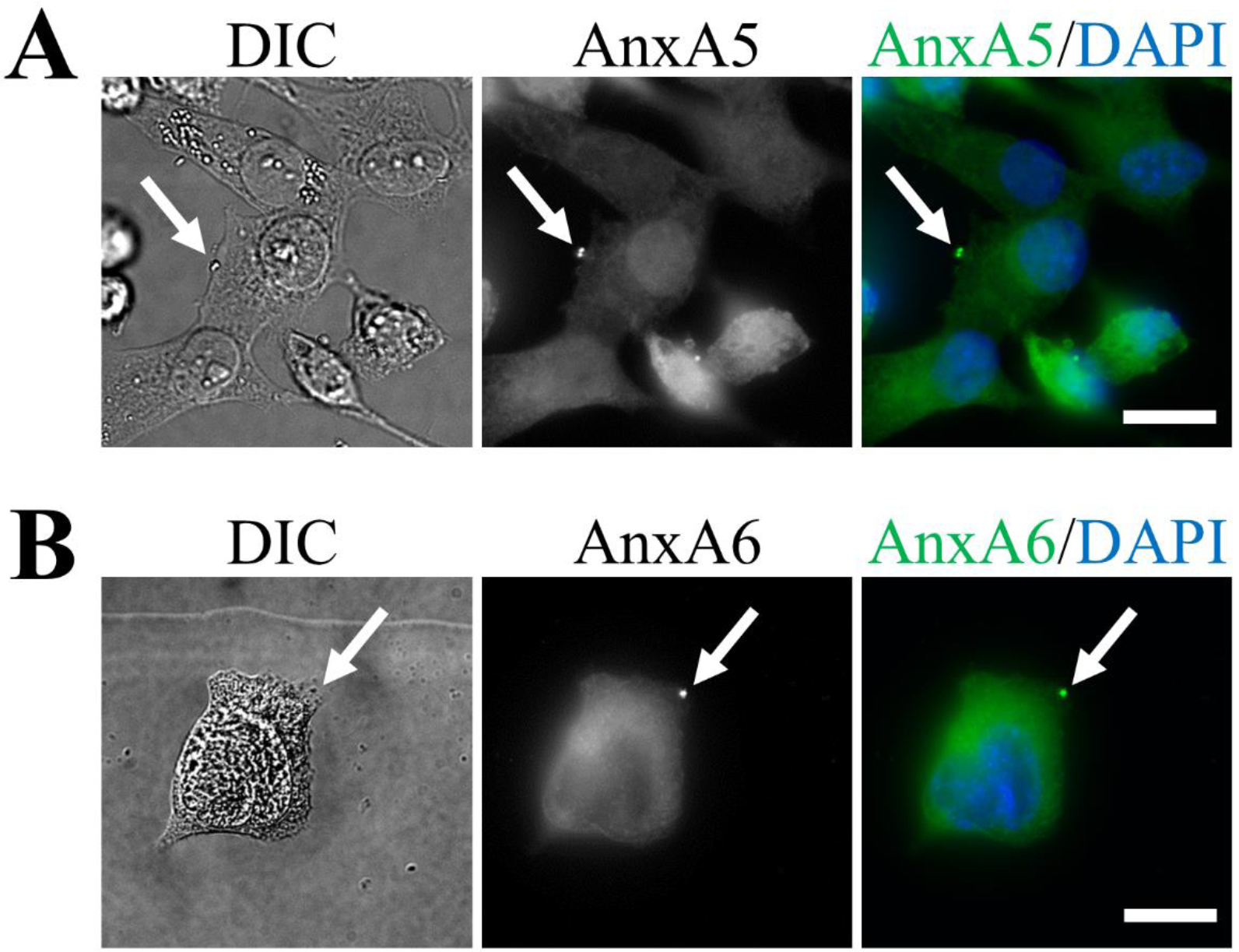
Subcellular localization of endogenous AnxA5 and AnxA6 in damaged MDA-MB-231. MDA-MB-231 cells were irradiated with a 110-mW infrared laser (white arrow) in DPBS+Ca2+, then fixed and immunostained for AnxA5 (**A**) or AnxA6 **(B)**. After laser injury, MDA-MB-231 cells exhibited an accumulation of AnxA5 or AnxA6 at the disruption site. Scale bar: 20 µm.

In order to confirm that AnxA5 and AnxA6 are required for membrane repair, both Anx were knocked-down in MDA-MB-231 cells using shRNA strategy. We estimated that the decrease of the expression of AnxA5 or AnxA6 was about 90% (**Fig. S2**). AnxA5-deficient or AnxA6-deficient MDA-MB-231 cells were then submitted to membrane repair assay. After laser injury, we observed that AnxA5-deficient MDA-MB-231 cells exhibited a rapid and strong increase in the intracellular fluorescence intensity due to the entry of FM1-43 dye, indicating a major defect of membrane repair (**Fig. 7, A and C, middle panel**). AnxA6-deficient MDA-MB-231 cells exhibited also a continuous entry of FM1-43 dye into the cytosol after membrane damage but in a lesser extent compared to AnxA5-deficient cells (**Fig. 7, B and C, right panel**). The absence of plateau within 120 s after membrane damage indicates that these cells present also a defect of membrane repair. 120 s after membrane injury, imaging of FM1-43 by video-microscopy was stopped and gain and offset values were adjusted for obtaining an unsaturated fluorescence image (**Fig. 7 D**). We observed that most AnxA5-deficient MDA-MB-231 cells (n = 45 /49) exhibited a large membrane disruption at the wound site (**Fig. 7 D, left panel**). We have proposed previously that AnxA5 molecules form 2D-arrays at the edges of the membrane disruption site in order to strengthen the membrane and to avoid the extension of the rupture (Bouter et al., 2011; Carmeille et al., 2015, 2016), which has been recently confirmed on model membrane (Lin et al., 2020). The hole present at the irradiation site suggests that plasma membrane has been torn. Lipid material, which is marked by the lipophilic dye FM1-43, is accumulated near the disruption site but looks insufficiently dense for plugging the disruption site. Strikingly, in the case of AnxA6-deficient MDA-MB-231 cells, we often (n = 36 /51) observed that lipid material is concentrated far away from the membrane injury (**Fig. 7 D, right panel)**. If the repair mechanism relies on the formation of a lipid “patch” that is supposed to clog the membrane disruption, AnxA6-deficient cells would suffer from a defect in the recruitment of this lipid “patch”. Altogether, these results led to conclude that AnxA5 and AnxA6 are instrumental for membrane repair in MDA-MB-231 cells.

**Figure 7.**
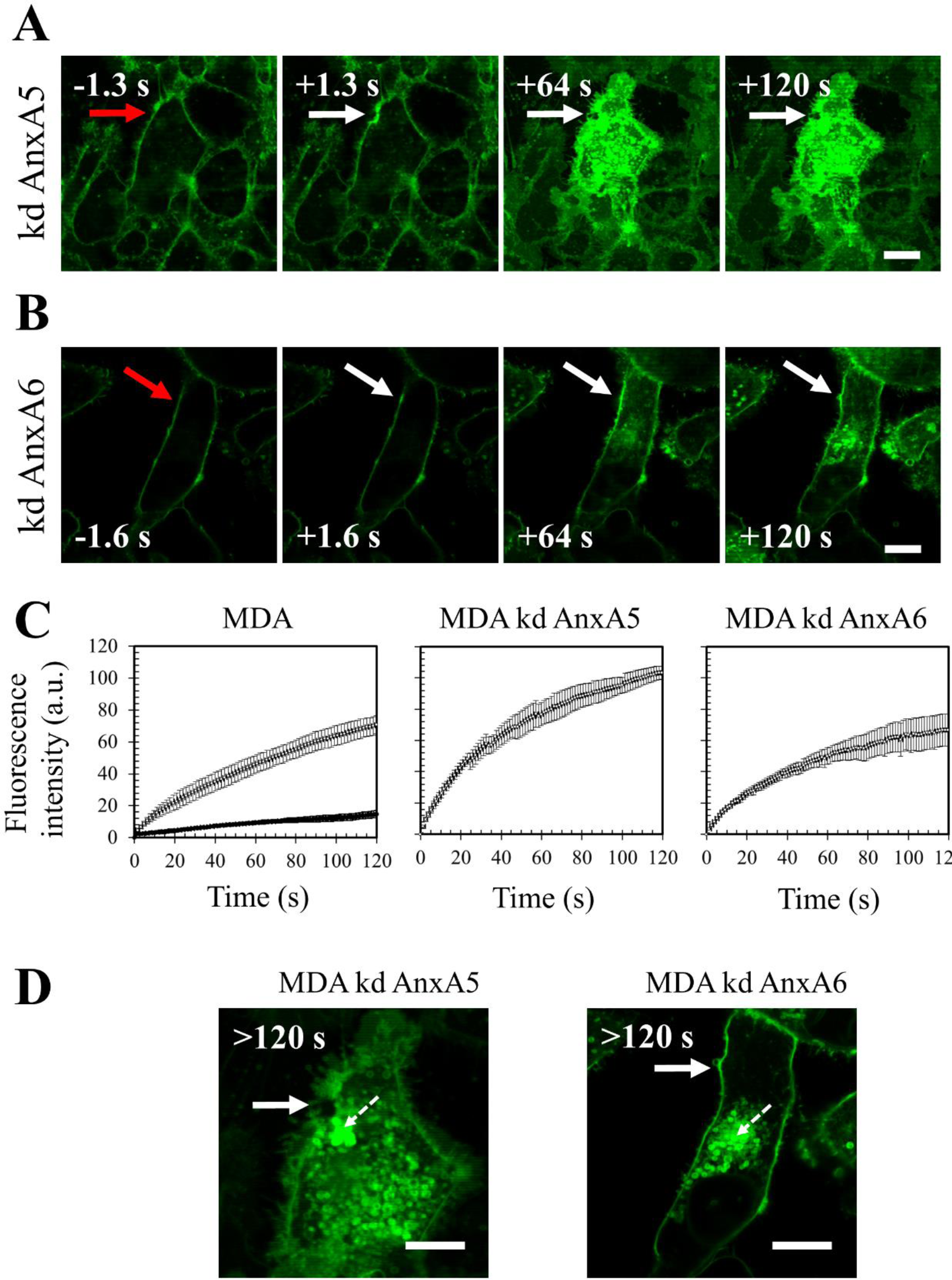
Responses of AnxA5-deficient or AnxA6-deficient MDA-MB-231 cells to a membrane damage by laser ablation. Sequence of representative images showing the response of an AnxA5-deficient **(A)** or an AnxA6-deficient MDA-MB-231. Scale bar: 10 μm. **(B)** cell after membrane damage performed by laser irradiation in the presence of FM1-43 (green) is presented. Cells were treated and presented as described in the legend of Figure 4. Representative images of about 50 cells. scale bar: 10 μm. **(C)** Kinetic data represent the FM1−43 fluorescence intensity integrated over whole cell sections, averaged for about 60 cells (+/−SD). For sake of clarity, data from the Figure **4C** concerning the responses of MDA-MB-231 cells (MDA) are presented. For AnxA5-deficient MDA-MB-231 (MDA kd AnxA5) or AnxA6-deficient MDA-MB-231 (MDA kd AnxA6) cells, a continuous and large increase of the fluorescence intensity was observed, indicating the absence of membrane resealing. **(D)** Zoomed and unsaturated images of kd AnxA5 and kd AnxA6 cells presented in (A) and (B) 120s after membrane injury, respectively. Large filled and small dashed white arrows point out the wound site and the lipid material accumulating inside the cell after membrane injury, respectively. Scale bar: 10 μm.

### AnxA5-AnxA6 deficient MDA-MB-231 cells suffer from a severe defect of membrane repair and die during migration on collagen I

We finally investigated the influence of knock-down in AnxA5 and/or AnxA6 on the migration of MDA-MB-231 cells on collagen I. For this purpose, we established a double knock-down AnxA5 and AnxA6 MDA-MB-231 cell line (hereafter named AnxA5-AnxA6 deficient MDA-MB-231 cells). MDA-MB-231 cells transduced together with AnxA5- and AnxA6-targeting shRNAs exhibited a decrease by about 73% and 99% for AnxA5 and AnxA6, respectively (**Fig. S3**). As expected, AnxA5-AnxA6 deficient MDA-MB-231 cells suffer from a defect of membrane repair after laser damage (**Fig. S4, A and B**). As observed for AnxA6-deficient MDA-MB-231 cells, lipid material appeared frequently concentrated far away from the membrane injury (**Fig. S4 C**).

In order to assess the influence of a defect of the membrane repair machinery in cell life during migration, we seeded 2.10^5^ control or Anx-deficient MDA-MB-231 cells on a dish coated with collagen I and cells were imaged and counted 24h and 48h after seeding. We hypothesized that migration of Anx-deficient MDA-MB-231 on collagen I may lead to the death of cells and their detachment from the coverslip, due to un-resealed membrane damages. 24h after seeding, we observed that Anx-deficient cells exhibit a cell density and morphology similar to control cells indicating that the deficiency in Anx does not disturb cell adhesion on collagen I-coated coverslip (**Fig. 8, 24h**). 48h after seeding, phase contrast microscopy imaging showed that cell density was similar between control and AnxA5-AnxA6 deficient cells, though the percentage of rounded cells was strongly increased (**Fig. 8, 48h**). Through video-microscopy analysis, we observed that elongated AnxA5-AnxA6 deficient MDA-MB-231 cells became rounded as soon as they were starting to migrate (**Video S6**). It was likely that rounded cells were dying cells, which were intended to be eliminated during washes prior to trypsinization. This was confirmed by cell counting, which showed that living adherent AnxA5-AnxA6 deficient cells represented in about half of the amount of control cells (**Fig. 8**). We observed that AnxA5 or AnxA6 deficiency had little effect on cell density and morphology, even the percentage of rounded cells appeared to be doubled for AnxA6-deficient MDA-MB-231 cells, without significant effect on the number of cells 48 h after seeding. A combined deficiency in AnxA5 and AnxA6 seems therefore to affect the migration of MDA-MB-231 cells on fibrillar collagen I.

**Figure 8.**
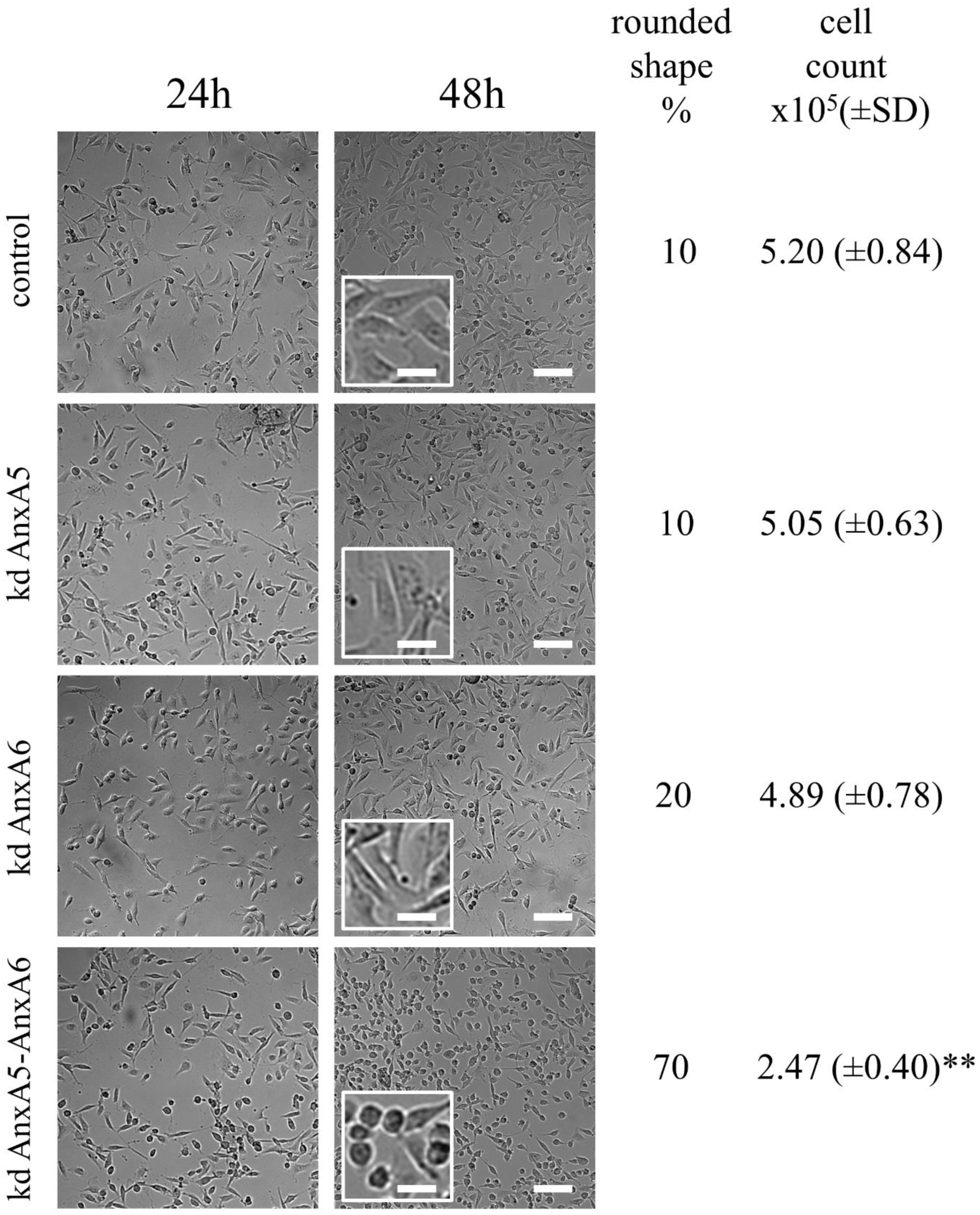
Responses of Anx-deficient MDA-MB-231 cells to membrane damages induced by migration on collagen I. 2.105 control, AnxA5-deficient, AnxA6-deficient or AnxA5-AnxA6 deficient MDA-MB-231 cells were seeded on a glass coverslip coated with collagen I. 24h (left panels) and 48h (middle panels) after seeding, cells were imaged by phase contrast microscopy. At 48h, the insert displays a magnified area from the image for sake of clarity. At 48h-post seeding, the percentage of cells displaying rounded shape was estimated using the ImageJ software. Rounded cells were defined as presenting a circularity ≥ 0.50, with circularity = 4*pi*(area/perimeter^2). Cells were then washed, trypsinized and counted. Mean values (± SD) from three independent experiments performed in triplicate are presented in the right lane. Student t Test for independent samples. **: p < 0.01. Scale bars: images, 100 μm; inserts, 25 µm.

To address the possibility that these AnxA5-AnxA6 deficient MDA-MB-231 cells suffer from a defect of membrane repair during their migration on collagen I, cells were loaded with Fluo-4-AM and kinetics study of cell migration on collagen I was performed by fluorescence microscopy. Images acquired in phase-contrast microscopy just before and after kinetics study ensured that cells have migrated (**Fig. 9 A**). We observed important variation of Fluo-4-AM fluorescence in several cells during migration (**Fig. 9 B)**. The strong increase of fluorescence intensity revealed the presence of rupture(s) in cell membrane as observed in control cells (**Fig. 3**). However, in the case of AnxA5-AnxA6 deficient MDA-MB-231 cells we observed that over a large period the fluorescence oscillated between a high intensity and a lesser one (**Fig. 9, B and C**). Through fluorescence video-microscopy, variations of intensity appeared as a flashing light (**Video S7**). It is unlikely that these variations were due to a series of damage and repair events. It is instead more relevant to think that it results from a strong increase of the intracellular calcium concentration subsequent to a membrane damage and an attempt from the cell to counteract this calcium excess, through calcium absorption by mitochondria for example. Altogether these results suggest that AnxA5-AnxA6 deficient MDA-MB-231 cells are unable to reseal plasma membrane damages induced by the migration on collagen I.

**Figure 9.**
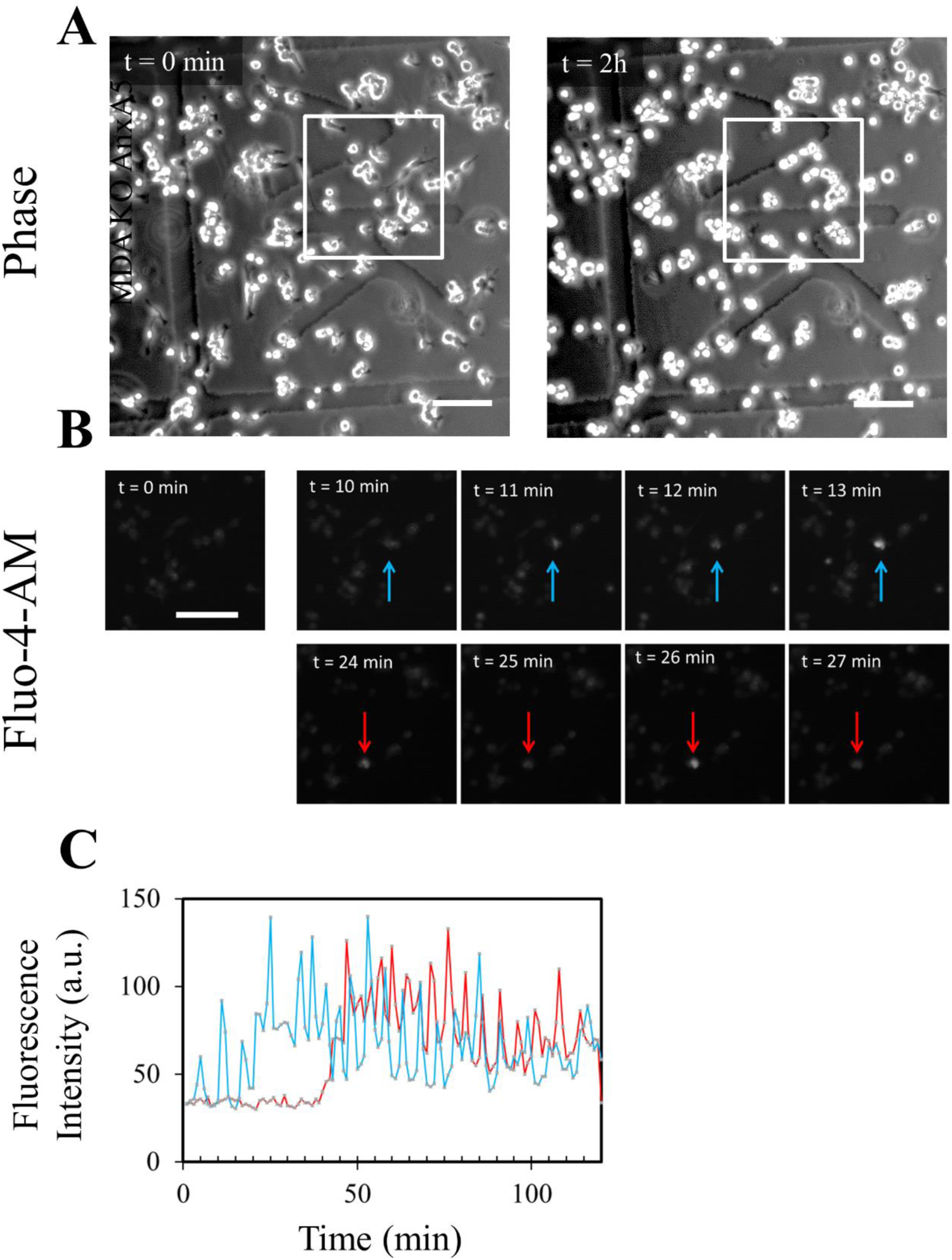
Highlighting un-resealed membrane damages in migrating AnxA5-AnxA6 deficient MDA-MB-231 cells. AnxA5-AnxA6 deficient MDA-MB-231 cells were treated as described in the legend of Figure 3. **(A)** Images in phase-contrast microscopy were acquired before and after the kinetics study performed by fluorescence microscopy. **(B)** Fluorescence intensity variations of intracellular Fluo-4-AM studied in the course of time. Each colored arrow points out a cell of interest exhibiting a strong variation of the fluorescence intensity. Strong increases of fluorescence intensity are particularly observed at t=11min and t=13min (blue arrow) and t=24min and t=27min (red arrow). A decrease of fluorescence intensity is observed at t=10min and t=12min (blue arrow) and t=25min and t=26min (red arrow). Scale bar: 100 µm. **(C)** Quantification of intracellular fluorescence intensity in arbitrary unit (a.u.) measured in cells presented in **(B)** during the two hours of migration. The color of each line corresponds to the color of the arrow in **(B)**.

## Discussion

### Migration on collagen I fibrils as a method for studying membrane repair

For reaching distant tissues, cancer cells are submitted to physical stresses during metastasis, due to their migration and invasion processes through dense extracellular matrix and due to intravasion and extravasion mechanisms for the entry into and the exit from the blood circulation, respectively. Shear stress induced by these processes may lead to cell membrane disruptions. This study aimed mainly at determining whether silencing of membrane repair machinery impairs survival of cancer cells during migration on extracellular matrix. It required first to find a method enabling to modelize shearing forces induced by the extracellular matrix. In this report, we have provided evidences that cell migration on collagen I fibrils leads to plasma membrane damages along with the release of membrane fragments in the wake of the cell. Membrane damage and release can be easily followed by fluorescence microscopy using labeled calcium indicator and plasma membrane stain. Beyond of its interest for research in cancer biology, this method is an outstanding tool for studying membrane repair in cells that exhibit the capacity of migration, notably epithelial and endothelial cells. It could replace current artificial methods such as laser irradiation, permeabilization using detergent, sprinkling with glass beads or scratching with a needle, that are far away from physiological injuries.

### AnxA5 and A6 belong to the membrane repair machinery of MDA-MB-231 cells

We then needed to identify proteins belonging to the membrane repair machinery. Several Anx have been shown to participate in membrane repair processes in different cell types (Jimenez and Perez, 2017; Draeger et al., 2011), and particularly in cancer cells (Sønder et al., 2019; Jaiswal et al., 2014; Boye et al., 2017). They constituted therefore prime targets for this achievement. Even though AnxA4 has been shown to play an instrumental role in membrane repair of MCF-7 cells (Boye et al., 2017), we observed that it is weakly expressed in MDA-MB-231 cells, suggesting its participation in membrane resealing is unlikely. However, AnxA1, A2, A5 and A6 are highly expressed in MDA-MB-231 cells and could be expected to participate in membrane resealing. Our attention focused particularly on AnxA5 and AnxA6, which exhibit varying intracellular concentrations. This heterogeneity of Anx expression probably explains discrepancy in the ability of MDA-MB-231 cells to reseal damaged plasma membrane. Our results demonstrate that AnxA5 and AnxA6 are instrumental for membrane repair in MDA-MB-231 cells. Whether the description of mechanisms related to membrane resealing in cancer cells was outside the scope of this study, some observations that we have done gave interesting clues on the role played by AnxA5 and AnxA6. We have previously proposed that AnxA5 forms 2D-arrays at the edges of the torn membrane allowing to strengthen the membrane and prevent the expansion of the tear (Bouter et al., 2011; Carmeille et al., 2015, 2016). This hypothesis has been recently confirmed by a sophisticated study performed by Scheuring and collaborators, using notably high-speed atomic force microscopy (Lin et al., 2020). They showed that AnxA5 self-assembly into lattices decreases lipid diffusivity, increases membrane thickness and increases lipid order. All together these results show that the formation of AnxA5 2D-arrays at the edges of the torn membrane induces a transition phase that stabilizes and structures the membrane into a gel phase. Here, we have shown that endogenous AnxA5 is recruited rapidly at the disruption site in MDA-MB-231 cells, where it probably self-assembles upon binding to the damaged membrane and promote membrane repair. Damage by laser irradiation in AnxA5-deficient MDA-MB-231 cells leads to the absence of membrane repair, which seems to be due to the expansion of the breach becoming too large to be plugged by the lipid “patch”. This observation is totally in line with the expected function of AnxA5 in membrane repair that we have previously reported.

AnxA6 is an unusual family member with two instead of one Anx core domains, each domain being composed by four 70-amino acid repeats. Together with other Anx, AnxA6 has been shown to assemble at the site of cell membrane damage and participate in membrane resealing in muscle (Swaggart et al., 2014; Roostalu and Strähle, 2012; Demonbreun et al., 2019) and cancer cells as well (Boye et al., 2017). A specific role for AnxA6 in membrane resealing of damaged MCF-7 cells have been previously proposed (Boye et al., 2017). From wound edges curved in out-of-plane by the action of AnxA4, AnxA6 induces constriction force responsible for the closure of the hole. It is important to note that MCF-7 cells were exposed to strong injuries by laser irradiation inducing a large wound diameter (up to 3 µm) in this study. As the membrane repair mechanism may vary depending on the cell type, the extent of the damage and the spatial position of the injury (Boye and Nylandsted, 2016; Boye et al., 2017; Jimenez and Perez, 2017), it is likely that the role played by AnxA6 for resealing MDA-MB-231 cells is slightly different in our study. Here, we show that AnxA6-deficient MDA-MB-231 cells suffer from a defect of membrane repair, which seems to be linked to the absence of the recruitment of cytoplasmic lipid material, ie the lipid “patch”, to the site of membrane damage. The presence of two Anx core domains gives the ability to AnxA6 to bridge two adjacent membranes, such as plasma membrane and cytoplasmic lipid vesicle. In addition, phylogenetic analysis of Anx has shown that the N-terminal core domain of AnxA6 is similar to AnxA3 whereas the C-terminal one is similar to AnxA5 (Morgan and Fernández, 1995). It has been shown that Anx induce different and specific membrane morphologies upon interaction (Boye et al., 2018). For instance, AnxA3 rolls the membrane in a fragmented manner from free edges, producing multiple thin rolls, whereas AnxA5 induces cooperative roll-up of the membrane. AnxA6 may therefore induces multiple membrane rearrangements at the disruption site at the interface between free edges of the damaged plasma membrane and the intracellular lipid “patch”. We hypothesize that the absence of AnxA6 may be responsible for the lack of recruitment of the lipid “patch”.

### Defective membrane repair machinery impairs survival of invasive cancer cells

The migration of MDA-MB-231 cells on fibrillar collagen I is accompanied by the release of membrane fragments responsible for the disruption of the cell membrane. In wild-type cells, in the presence of AnxA5 and AnxA6, these membrane injuries are resealed in less than a minute (**Fig. 10 A**). As observed for damage by laser irradiation, it is likely that the increase of intracellular calcium concentration induces the recruitment of Anx at the disruption site, where they promote membrane resealing. Once membrane repair is done, Anx come back to the cytosol and the cell can continue to migrate. The combined inhibition of the expression of AnxA5 and AnxA6 leads to a major defect of the membrane repair machinery. In MDA-MB-231 cells, this silencing leads to the incapacity of cells to migrate on fibrillar collagen I (**Fig. 10 B**). In this case, disruptions due to migration are indeed not repaired leading to cell death. Consequently, cell migration is finally discontinued. Strikingly, we have observed that the inhibition of only AnxA5 or AnxA6 was not sufficient to induce death of cells and the stop of migration. It is likely that membrane ruptures created by shear stress on collagen fibrils were minor forces leading to small ruptures (few hundreds nm), smaller than those created by laser irradiation (1 µm). In this case, the low residual concentration of AnxA5 or AnxA6 may be sufficient for the achievement of membrane repair. One can also envision that the lack of AnxA5 or AnxA6 may be compensated by another Anx. In a physiological context, we hypothesize that mechanical stress would be stronger than in our *in vitro* experimental conditions. Indeed, a cell that migrates *in vivo* through the extracellular matrix is submitted to shear forces on the whole surface of the plasma membrane, whereas in our *in vitro* experimental model shear forces affect only plasma membrane at the interface interacting with the substrate. In addition, we can expect that shear forces induced by other cells during intravasion and extravasion processes are likely to be forces with greater strength.

**Figure 10.**
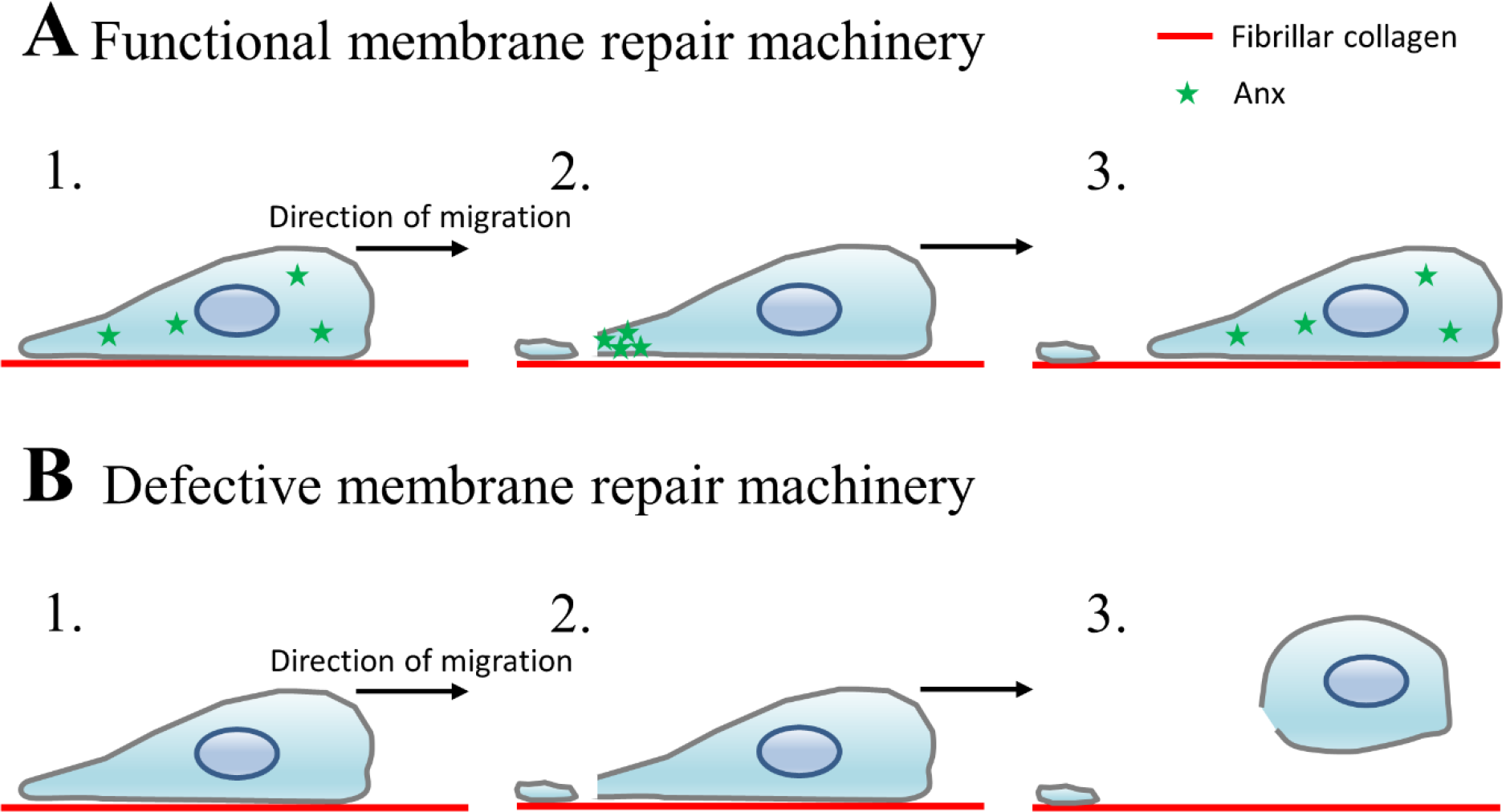
Model of migration of MDA-MB-231 cells equipped with a functional or defective membrane repair machinery. **(A)** In wild-type cells, migration on fibrillar collagen I induces the release of membrane fragments responsible for membrane injuries (Step 1). Anx, notably AnxA5 and AnxA6, are recruited to the disruption site, where they promote membrane repair (Step 2). Once the cell is repaired, Anx return to the cytosol (Step 3). **(B)** In cells presenting defective membrane repair, due for example to the absence of Anx, membrane injuries created by the migration on collagen I fibrils are not resealed (Step 2). This leads to the death of cells, which are released from the substrate (Step 3).

In conclusion, we show here that silencing of the membrane repair machinery induces a stop of migration of cancer cells and constitutes a promising approach for inhibiting cancer metastasis.

## Materials and methods

Cell culture media and reagents were from ThermoFisher Scientific (Waltham, MA, USA) except when otherwise stated.

### Cell culture

MDA-MB-231 cells were cultured in Dulbecco modified Eagle’s minimal essential medium (DMEM) containing 4 mM Glutamax© and supplemented with 10% fetal bovine serum and penicillin / streptomycin (100 U/mL and 100 μg/mL). The choriocarcinoma BeWo cell line was cultured in Ham’s F12K medium containing 5 mM L-glutamine and supplemented with 20% fetal bovine serum and penicillin / streptomycin (100 U/mL and 100 μg/mL). Both cell types were maintained at 37°C in a 5% CO2 humidified incubator.

### Coating

All experiments were performed in 35-mm glass bottom dishes equipped with a square patterned coverslip (MatTek, Ashland, USA). Coating was performed as previously described(Di Martino et al., 2017). Briefly, the glass coverslip was first coated for 20 min at RT with 1 mg/mL gelatin, which was fixed by 0.5% glutaraldehyde for 40 min at RT. After three washes with Dulbecco’s phosphate buffer saline (DPBS) containing 0.9 mM CaCl2 (DPBS+Ca^2+^), collagen I at 0.5 mg/mL was incubated for 4 h at 37°C. After removing the excess of collagen I, 4.10^4^ cells in growth medium were subsequently seeded and cultured for 24 h.

### Kinetics study of MDA-MB-231 migration by phase contrast video-microscopy

Phase-contrast video-microscopy was performed using the Leica DMI6000B microscope equipped with CCD camera (6.45 μm pixel dimension, 1392 x 1040 pixels, Leica) 48 h after cell seeding. Dishes were positioned in the tempered (37°C) humidified chamber at least 30 min before beginning kinetics study. Unless otherwise indicated, kinetics studies lasted 2 h using an acquisition interval of 1 min. Tracking of cells was performed using the MTrackJ pluging from ImageJ software (Meijering et al., 2012). For each condition, at least 100 cells from three independent experiments were analyzed.

### Kinetics study of Fluo4AM-loaded MDA-MB-231 migration by fluorescence microscopy

Working solution was prepared by dissolving 50 µg of Fluo-4AM in DMSO at a final concentration of 1 mM. 10 µL of Fluo-4AM working solution were mixed with 8 µL of Pluronic F-127 (Sigma). The mixture was vortexed and diluted in 2 mL growth medium, which were added on cells for 1 h at 37°C. After one wash with DPBS+Ca^2+^, cells were incubated in growth medium. Fluorescence and phase-contrast video-microscopy was performed using the Leica DMI6000B microscope. Kinetics studies lasted 2 h with 30-s interval acquisitions.

### CellMask™Orange staining

A fresh working solution was prepared by diluting the commercial CellMask™Orange stock solution at 1:1000 in 2 mL DPBS+Ca^2+^. After 2 h of migration on collagen I, the growth medium of MDA-MB-231 cells was replaced by the CellMask™Orange working solution for 5 min incubation at 37°C. Cells were washed three times in DPBS+Ca^2+^ solution and then fixed in PFA4% for 10 min at RT. After three washes in DPBS+Ca^2+^, CellMask™Orange staining was observed using the cube U-MWG2 containing in a band-pass excitation filter (510-550 nm), a dichroic mirror (50 % transmission from 570 nm) and a long-pass emission filter (threshold 590 nm) from the conventional fluorescence IX81 microscope (Olympus). Images were analyzed with the ImageJ software.

### Membrane rupture and repair assay

Membrane repair assay differed slightly from our previous studies (Carmeille et al., 2017). MDA-MB-231 cells were cultured in complete growth medium on 18*18 mm glass coverslips (Nunc). A solution of 5 µg/mL FM1-43 (ThermoFisher Scientific, Waltham, MA, USA) in D-PBS containing 2 mM Ca^2+^ was maintained over ice and subsequently added in a homemade coverslip cell chamber (**Fig. S5**), where cells containing coverslip was mounted. To induce membrane damage, cells were irradiated at 820 nm with a tunable pulsed depletion laser Mai Tai HP (Spectra-Physics, Irvine, USA) of an upright two-photon confocal scanning microscope (TCS SP5, Leica) equipped with an HCX PL APO CS 63.0 x 1.40 oil-objective lens. Irradiation consisted of 1 scan (1.3 s) of a 1 µm x 1 µm area with a power of 110 (±5) mW. 512 x 512 images were acquired at 1.3 s intervals with pinhole set to 1 Airy unit. Membrane rupture and repair processes were monitored by measuring variations in fluorescence intensity of FM1-43 as previously described (Carmeille et al., 2017). FM1-43 was excited by the 488-nm laser line (intensity set at 30% of maximal power) and fluorescence emission was measured between 520 nm and 650 nm. For quantitative analysis, the fluorescence intensity was integrated over the whole cell surface and corrected for the fluorescence value recorded before irradiation, using ImageJ software.

### Western-blot

2.10^6^ cells were trypsinized, pelleted and re-suspended in 300 µL of D-PBS supplemented with 1 mM EGTA. Protein extracts were obtained by sonicating ice-cold cell suspension with a Branson digital sonifier (amplitude 20%, duration 2 min, interval 5 s and pulse 5 s). Two successive centrifugations at 13,000g for 1 min allowed to remove cell debris. 10 µg protein extracts were separated on a 10% SDS-PAGE for detection of Anx. Semi-dry electrophoretic transfer (Bio-Rad, Hercules, CA, USA) onto PVDF membrane was performed for 1 h at 100 V. The cellular content in AnxA1 (37 kDa), AnxA2 (36 kDa), AnxA4 (32 kDa), AnxA5 (35 kDa), AnxA6 (68 kDa), and anti-glyceraldehyde-3-phosphate deshydrogenase (GAPDH, loading control, 37 kDa) was detected with rabbit anti-AnxA1 polyclonal antibody (PA1006, BosterBio, Pleasanton, CA, USA), mouse anti-AnxA2 monoclonal antibody (3E8-B6, Sigma, Saint-Louis, MO, USA), mouse anti-AnxA4 monoclonal antibody (SAB4200121, Sigma), anti-AnxA5 monoclonal antibody (AN5, Sigma), mouse anti-AnxA6 monoclonal antibody (sc-271859, Santa cruz Biotechnology), and rabbit anti-GAPDH polyclonal antibody (FL-335, Santa Cruz Biotechnology), respectively. All primary antibodies were diluted 1:1,000 in saturation solution composed by Tris buffer saline (10 mM Tris, 150 mM NaCl, pH 8.0) supplemented with 0.1% Tween20 and 5% non-fat dry milk. Revelation was performed using secondary antibodies conjugated to horse-radish peroxidase (GE-Healthcare) diluted 1:2,000 in saturation solution and Opti-4CN™ colorimetric kit (Bio-Rad). ImageJ software was used to measure the relative intensity of protein bands.

### Transduction of AnxA5-targetting and/or AnxA6-targetting shRNA lentiviral particles in MDA-MB-231 cells

The following shRNA sequences, cloned into the pLKO.1 puro-vector (MISSION® shRNA plasmids, Sigma), were used: AnxA5-targetting shRNA (A5shRNA): 5’-CCGGCGCGACTT CTGGAATTTACTCGAGTAAATTGCCAGAAGTCTCGCGTTTTT-3’; AnxA6-targetting shRNA (A6shRNA): 5’-CCGGCGGGCACTTCTGCCAAGAAATCTCGAGATTTCTTGGCAGAAGTGCCCGTTTTT-3’; scrambled shRNA (SCshRNA): 5’-CCTAAGGTTAAGTCGCCCTCGCTCGAGCGAGGGCGACTTAACCTTAGG-3’.

Lentiviral-based particles containing shRNAs were produced by Bordeaux University Lentiviral Vectorology Platform (US005, Bordeaux, France) by transient transfection of 293T cells. 2.10^5^ MDA-MB-231 cells were cultured in a 30 mm Petri Dish for 24 h and transduction was carried out by adding concentrated lentiviral particles to the cells at multiplicity of infection (MOI) of 10 in 2 mL Opti-MEM® for 24 h. Transduced cells were cultured for 24 h in growth medium and then selected in selection medium composed by 2 µg/mL puromycin in DMEM for 48 h. Cells were passed and subsequently cultured in 25 cm^2^ cell culture flask in selection medium. At each passage, a fraction of cells was used for preparing protein extracts for western-blot analysis of the expression of endogenous Anx. Co-transduction of lentiviral-based particles containing AnxA5- and AnxA6-targetting shRNAs was performed by mixing each type of particles at MOI of 10 with 2.10^5^ MDA-MB-231 cells, which corresponds to a final MOI of 20.

### Localization of endogenous Anx in intact and damaged cells by immunofluorescence

For subcellular localization of endogenous Anx in damaged MDA-MB-231 cells, cell membrane rupture was performed according to the protocol described above, but in the absence of FM1-43 to avoid fluorescence cross-talk. After laser irradiations, cells were fixed in 1% glutaraldehyde and permeabilized in 0.1% TritonX100 diluted in DPBS+Ca^2+^. All subsequent steps (saturation, antibody incubation and washes) were performed using 2% BSA in DPBS+Ca^2+^ solution. Primary antibodies (1:100, except for anti-AnxA5 1:400), which were the same as used for western-blot, and secondary Alexa Fluor 488-coupled anti-mouse or anti-rabbit IgG goat antibodies (1:1000, ThermoFisher Scientific) were successively incubated with cells for 1 h at 37°C. Finally, cells were washed in DPBS+Ca^2+^ and nuclear counterstaining was performed with DAPI (Sigma). For each condition, about 30 cells from three independent experiments were analyzed.

For the subcellular localization of endogenous Anx in intact MDA-MB-231 cells, cells were cultured in 8-well Lab-Tek™ chambered coverglasses. Cells were immunostained according to the protocol described above for the localization of Anx in damaged cells, from the step of glutaraldehyde fixation.

### Online supplemental material

**Fig. S1** shows that MDA-MB-231 cells that express AnxA5 at a high level express also strongly AnxA6 and, inversely, a low level of AnxA5 is systematically accompanied by a weak expression of AnxA6. **Fig. S2** shows that the decrease of the expression of AnxA5 or AnxA6 in knocked-down MDA-MB-231 cells is about 90%. **Fig. S2** shows also specificity of action of A5shRNA and A6shRNA sequences in knock-down experiments. **Fig. S3** presents western-blot experiment showing that MDA-MB-231 cells transduced together with AnxA5- and AnxA6-targeting shRNAs exhibit a decrease by about 73% and 99% for AnxA5 and AnxA6, respectively. **Fig. S4** shows the response of AnxA5-ANxA6 deficient MDA-MB-231 cells to a membrane damage by laser ablation, which leads to conclude that these cells suffer from a defect of membrane repair. **Fig. S5** presents the cell culture device used for membrane repair assay. **Video S1 and S2** present the migration of MDA-MB-231 cells respectively in the presence and in the absence of collagen I using phase-contrast video-microscopy. **Video S3** presents a cell of interest that have migrated on fibrillar collagen I, for which CellMask™staining revealed the presence of membrane fragments in its wake. **Video S4** shows the important variations of fluorescence intensity in Fluo-4-AM loaded MDA-MB-231 cells migrating on collagen I using fluorescence video-microscopy. **Video S5** shows the responses of MDA-MB-231 cells to a membrane damage by laser ablation. **Video S6** shows that AnxA5-AnxA6 deficient MDA-MB-231 cells become rounded as soon as they are starting to migrate in collagen I, using phase-contrast video-microscopy. **Video S7** shows by fluorescence video-microscopy that Fluo-4-AM loaded AnxA5-AnxA6 deficient MDA-MB-231 cells exhibits repeated variations of the intracellular fluorescence.

## Abbreviations

Anx: Annexins;
AnxA1: Annexin-A1;
AnxA2: Annexin-A2;
AnxA5: Annexin-A5;
AnxA6: Annexin-A6;
Ca^2+^: calcium;
DPBS: Dulbecco’s phosphate buffer saline;
MOI: multiplicity of infection;
PS: Phosphatidylserine.

## Acknowledgments

This work was funded by the Cancéropole Grand-Sud-Ouest (grant 2015-E21 to A.B.) and Ligue Contre Le Cancer Gironde (grant 191796 to A.B.). The help of Marion Decossas, Elise Dargelos, Sylvie Poussard and Olivier Lambert is acknowledged for the use the Leica DMI6000B microscope. Christel Poujol and Sébastien Marais are acknowledged for the help in membrane repair assays that were done in the Bordeaux Imaging Center a service unit of the CNRS-INSERM and Bordeaux University, member of the national infrastructure France BioImaging supported by the French National Research Agency (ANR-10-INBS-04). The authors thank the staff of Vect’UB, the vectorology platforms (INSERM US 005 – CNRS UMS 3427-TBM-Core, Université de Bordeaux, France) for technical assistance. The authors declare no competing financial interests.

## Author contributions

F. Bouvet and A. Bouter performed the majority of experiments with assistance from M. Ros (study of migrasome and set-up of protocol for kinetics study of cell migration), E. Bonedeau (study of migrasome), C. Croissant (analysis by western-blot of Anx-deficient cells), L. Frelin (analysis of survival of Anx5- or AnxA6-deficient cells). A. Bouter coordinated the entire project. A. Bouter, V. Moreau and F. Saltel designed the experiments and contributed to the writing of the manuscript.

