## Supplemental material Fig S1-S5 for "Defective membrane repair machinery impairs survival of invasive cancer cells"

**Figure S1**

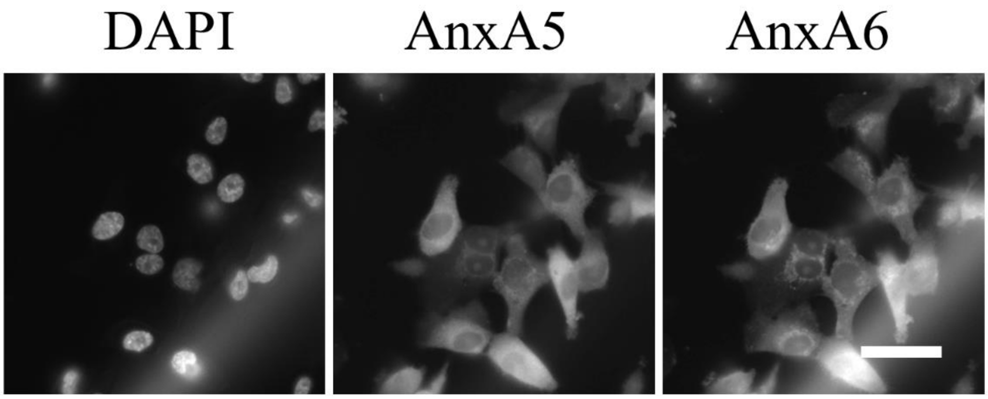


**Co-immunostaining of endogenous AnxA5 and AnxA6 in MDA-MB-231 cells.** MDA-MB-231 were jointly immunostained for AnxA5 and AnxA6 as described in Figure 5. Primary antibodies were rabbit polyclonal anti-AnxA5 and mouse monoclonal anti-AnxA6 and secondary antibodies were Alexa546-conjugated goat anti-rabbit IgG and Alexa488-conjugated goat anti-mouse IgG, respectively. MDA-MB-231 cells that strongly express AnxA5 present also a high level of AnxA6 expression and inversely. Scale bar: 40 µm.

**Figure S2**

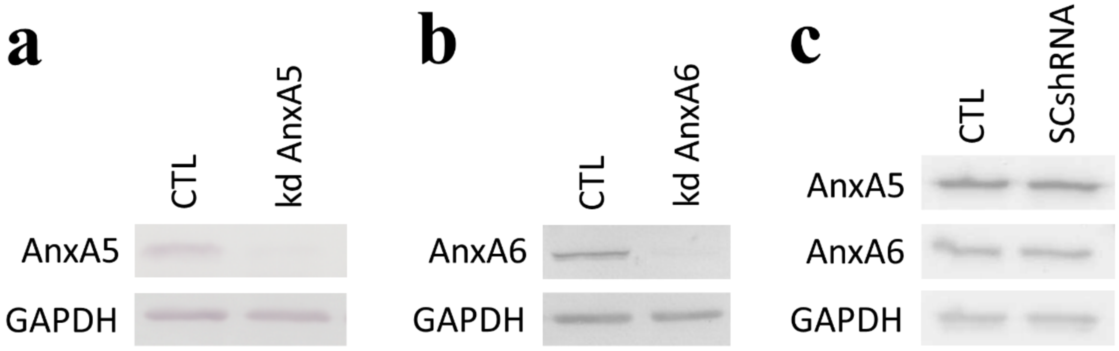


**Knock-down of AnxA5 or AnxA6 in MDA-MB-231 cells by shRNA strategy.** MDA-MB-231 cells were transduced (MOI =10) with lentiviral particles containing A5shRNA targeting AnxA5 (**a**), A6shRNA targeting AnxA6 (**b**) and SCshRNA (**c**), a scrambled shRNA. Non-transduced cells (CTL) were used as control. The cellular content of AnxA5 or AnxA6 was quantified by western-blot. The equivalent of 10 µg of protein extracts was separated by SDS-PAGE with 10% polyacrylamide. AnxA5 or AnxA6 were detected with primary mouse monoclonal antibodies and GAPDH (loading control) was detected with primary rabbit polyclonal antibody. The expression of AnxA5 or AnxA6 is decreased of more than 90% in A5shRNA or A6shRNA transduced MDA-MB-231 cells, respectively. No difference was observed between MDA-MB-231 cells transduced with SCshRNA and non-transduced cells regarding the expression of AnxA5 and AnxA6, which ensures the specificity of A5shRNA and A6shRNA sequences.

**Figure S3**

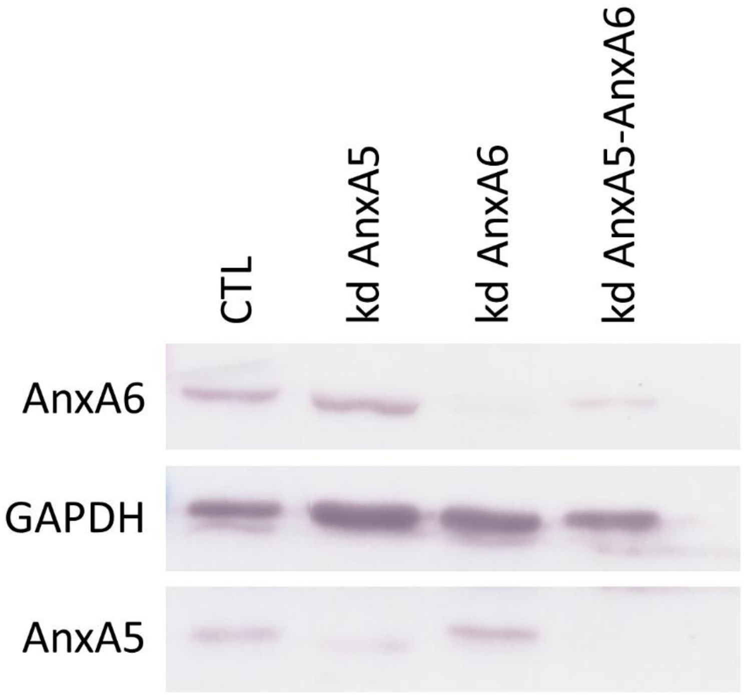


**Knock-down of AnxA5 and AnxA6 in MDA-MB-231 cells by shRNA strategy.** MDA-MB-231 cells were transduced (MOI =20) with lentiviral particles containing A5shRNA targeting AnxA5 and A6shRNA targeting AnxA6 for generating AnxA5-AnxA6 deficient cells (kd AnxA5 + kd AnxA6). The cellular content of AnxA5 and AnxA6 was quantified by western-blot in these cells and compared to kd AnxA5, kd AnxA6 and non-transduced (CTL) MDA-MB-231 cells. Western-blot was performed as described in the legend of supplementary Figure 2. In AnxA5-AnxA6 deficient MDA-MB-231 cells, the expression of AnxA5 and AnxA6 is decreased of about 73% and 99%, respectively.

**Figure S4**

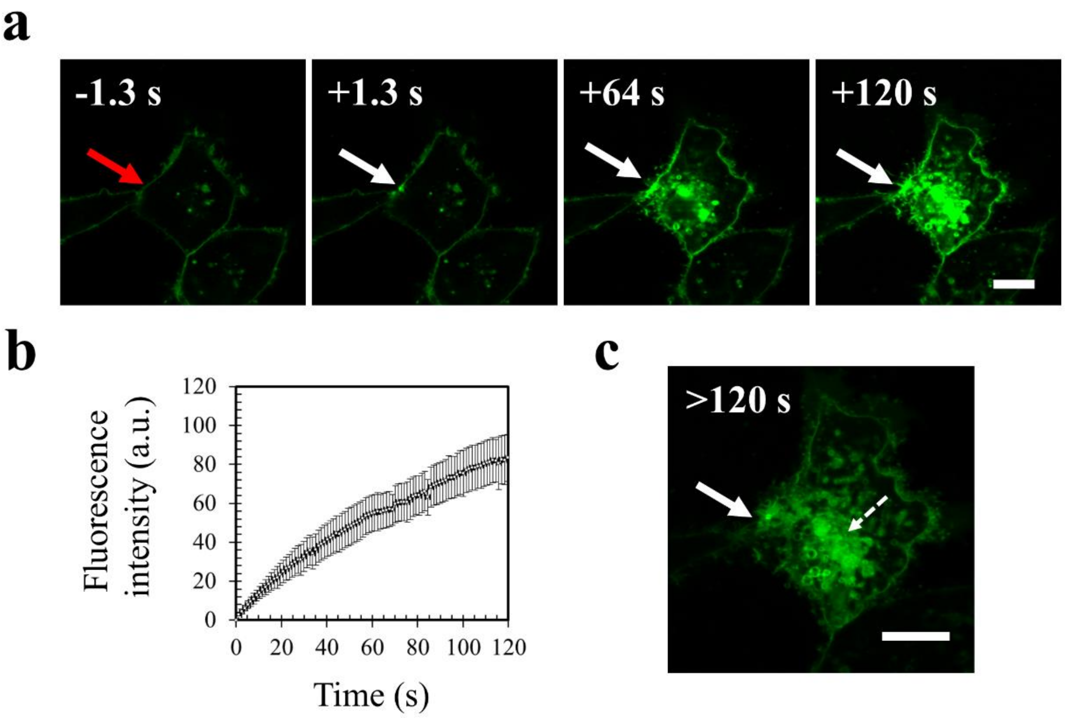


**Response of AnxA5-AnxA6 deficient MDA-MB-231 cells to a membrane damage by laser ablation.** MDA-MB-231 cells were rendered deficient for AnxA5 and AnxA6 by transduction with specific shRNAs (see Supplementary Figure 3). **a** Sequence of representative images showing the response of an AnxA5-AnxA6 deficient MDA-MB-231 cell after membrane damage performed by laser irradiation in the presence of FM1-43 (green). Cells were treated and presented as described in the legend of Figure 4. **b** Kinetic data represent the FM1−43 fluorescence intensity integrated over whole cell sections, averaged for about 60 cells (+/−SD). A continuous and large increase of the fluorescence intensity was observed, indicating the absence of membrane resealing. **c** Zoomed and unsaturated images of AnxA5-AnxA6 deficient cell presented in (**a**) 120s after membrane injury. Large filled and small dashed white arrows point out the wound site and the lipid material accumulating inside the cell after membrane injury, respectively. Scale bars: **a and c,** 10 µm.

**Figure S5**

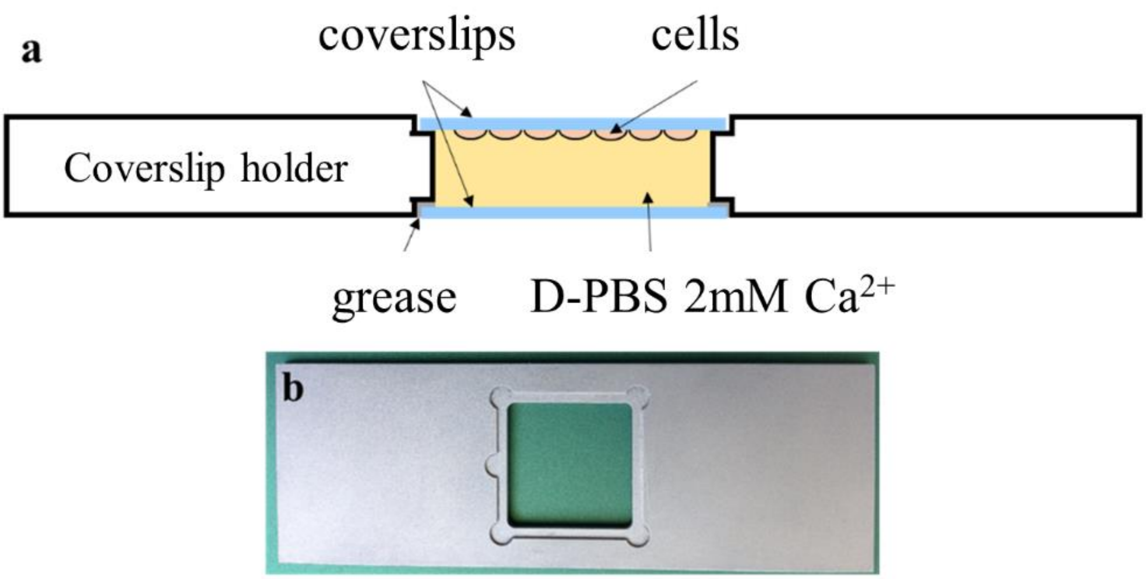


**Chambered cell culture slide used for cell membrane damage by laser irradiation with the upright two-photon confocal scanning microscope TCS SP5 (Leica). a** Schematic representation of the assembly with the coverslip holder and the coverslip on which cells have been cultured. **b** Photography of the metal coverslip holder for 18x18 mm coverslips.

**Supplemental videos**

**Video S1: Phase-contrast video-microscopy of the migration of MDA-MB-231 cells in the presence of collagen I.** See the legend of Fig. 1 A. Compared to Fig. 1 a larger field is presented (field width = 600µm). Frame rate = 10 fps.

**Video S2: Phase-contrast video-microscopy of the migration of MDA-MB-231 cells in the absence of collagen I.** See the legend of Fig. 1 A. Compared to Fig. 1 a larger field is presented (field width = 550µm). Frame rate = 10 fps.

**Video S3: Phase-contrast video-microscopy of a migrating MDA-MB-231 cells on collagen I.** See the legend of Fig. 2D. Field width = 360 µm. Frame rate = 20 fps.

**Video S4: Fluorescence video-microscopy of Fluo-4-AM loaded MDA-MB-231 cells on collagen I.** See the legend of Fig. 3 B. Field width = 250 µm. Frame rate = 10 fps.

**Video S5: Response of MDA-MB-231 cells to a membrane damage by laser ablation.** See the legend of Fig. 4. Video S5 A and B show the response of a MDA-MB-231 resealing (a) or not (b) a membrane damage performed by laser irradiation, respectively. Field width = 90 µm. Frame rate = 5 fps.

**Video S6: Phase-contrast video-microscopy of AnxA5-AnxA6 deficient MDA-MB-231 cells on collagen I.** See the legend of Fig. 10. Field width = 150 µm. Frame rate = 10 fps.

**Video S7: Fluorescence video-microscopy of Fluo-4-AM loaded AnxA5-AnxA6 deficient MDA-MB-231 cells on collagen I.** See the legend of Fig. 10 B. Field width = 150 µm. Frame rate = 10 fps.
